# CPMsi: a data-driven Computational Pipeline for fast, large-scale Mass Spectrometry Imaging analysis

**DOI:** 10.64898/2026.10.15.758756

**Authors:** Paola Tasca, Iulia Sidorov, Hans J. Dalebout, Nina Ogrinc, Ron M. A. Heeren, Aiko P. de Vries, Jesper Kers, Manfred Wuhrer, Ton J. Rabelink, Martin Giera, Yassene Mohammed

## Abstract

Mass Spectrometry Imaging (MSI) is increasingly used in clinical research to characterize tissue heterogeneity and identify disease-associated molecular patterns while preserving spatial context. However, translating MSI to large patient cohort studies remains challenging because, among others, existing computational tools struggle to scale with the size and complexity of such datasets. To overcome this bottleneck, we developed CPMsi, a scalable, data-driven computational pipeline for cohort-scale MSI analysis. CPMsi directly processes raw MSI data, performs global spectral alignment using a novel binning strategy, identifies regions of molecular similarity, and generates consensus spectra that compactly capture spatial molecular heterogeneity. The pipeline supports mapping of regions across consecutive tissue sections, and integration of multi-modal and multi-omics MSI acquisitions from the same section. We tested CPMsi using (i) consecutive brain MSI sections comprising ∼600,000 and ∼500,000 pixels with ∼20,000 *m/z* features, (ii) multimodal brain MSI lipidomic acquired from the same tissue section in positive and negative ionization modes, and (iii) a multi-omics brain MSI dataset of lipidomic, metabolomic, and glycomic measurements. CPMsi identified regions of molecular similarity 30-times faster than vendor-provided MSI analysis software. Consensus spectra substantially reduced data complexity while preserving molecular differences between tissue regions and enabled mapping regions of similar molecular composition across consecutive tissue sections. Integration of complementary MSI modalities further revealed molecular patterns that were not resolved by individual modalities alone. Overall, CPMsi enables efficient reduction, comparison, and integration of complex MSI datasets, providing a scalable framework for cohort-scale studies, biomarker discovery, and future clinical applications.

## Introduction

Mass spectrometry imaging (MSI) is an analytical technique that enables simultaneous measurement of the spatial distribution of thousands of molecules such as metabolites, lipids, and proteins, directly on tissue sections. By combining spatially resolved molecular characterization with histological context, MSI can reveal disease-specific molecular patterns that are not visible with conventional approaches ^1, 2, 3, 4, 5.^. Translating these insights into clinically applicable biomarkers and diagnostic tools requires robust validation in large patient cohorts and multicenter settings ^6^. However, scaling MSI studies to this level increases data volume and analytical complexity, making robust comparisons across samples and studies challenging ^7^.

A wide range of computational methods have been developed to optimize specific processing tasks, including mass recalibration ^8^, spectral alignment ^9^, and peak detection ^10^. While these methods improve individual analysis steps, they are often distributed as standalone algorithms and therefore require users to assemble complex analysis workflows from multiple tools.

Integrated software solutions exist, with SCiLS Lab (Bruker Daltonics) being one of the most widely used platforms for visualization, preprocessing, segmentation, and statistical analysis. Nevertheless, its application to comprehensive end-to-end analysis of large MSI cohorts remains challenging. Furthermore, its workflow requires substantial user intervention, e.g. adjusting peak width and shape, setting signal-to-noise thresholds, which can limit scalability and hinder efficient analysis of large cohorts.

On the other hand, open-source end-to-end frameworks have been developed to overcome some of these limitations. Examples include pyBASIS which enables robust comparisons across large tissue collections through alignment of chemically related ions ^11^; Cardinal which provides memory-efficient approaches for large multi-tissue experiments ^12^; M2aia for fast interactive visualization and processing of large 2D/3D MSI datasets ^13^; uMAIA which implements an unsupervised strategy to automatically match peaks across acquisitions ^14^; and msiFlow which offers an automated workflow for high-throughput and multimodal MSI analysis ^15^. These tools represent an important step toward more accessible and scalable MSI data analysis and highlight the potential of workflow-based approaches for processing large and complex datasets. However, many of these workflows are user-driven, i.e. to some level, they remain dependent on user-defined parameter choices and often employ dimensionality reduction or feature-selection strategies to reduce data complexity. While these approaches can improve computational efficiency, they emphasize dominant sources of variation while potentially overlooking low-abundance or subtle molecular signals which might be biologically relevant.

To overcome these limitations, we introduce CPMsi, a scalable, data-driven computational pipeline for the analysis of large-scale MSI studies. CPMsi directly processes raw spectra and aligns their *m/z* values using a novel binning approach, enabling spectrum-to-spectrum similarity calculations that are more than 100 times faster than up-to-date methods. These similarity measures support multiple downstream applications, including hierarchical clustering to group pixels into molecular regions, comparison of spatial patterns, and integration across datasets. CPMsi also generates consensus spectra for the identified regions, providing a compact representation that preserves relevant spectral features while substantially reducing data complexity. This enables efficient identification of region-specific molecular signatures and robust comparisons across tissues and patients, thereby facilitating MSI analysis at cohort scale.

We applied CPMsi to high-resolution MSI lipidomics data from two consecutive mouse brain sections comprising more than 600,000 and 500,000 pixels, respectively, across an *m/z* range of 260-1,100 to evaluate its computational efficiency and scalability. To assess its data integration capabilities, we further analyzed sequential multimodal lipidomic measurements acquired in positive and negative ion modes. In addition, we applied CPMsi to a publicly available multi-omics dataset comprising metabolomic, lipidomic, and glycan measurements from the same tissue section. Together, these datasets allowed us to evaluate CPMsi across different spatial resolutions, ionization modes, and molecular modalities, demonstrating its ability to efficiently process high-dimensional MSI datasets and highlighting its potential for larger-scale MSI studies.

## Results

CPMsi is a python library comprising four modules (Figure 1) that can be executed end-to-end, or independently, facilitating re-analysis of specific processing steps without re-running the entire workflow. CPMsi differs from traditional MSI analysis workflows by shifting from a peak-to-region approach to a region-to-peak strategy. Rather than selecting molecular features before spatial analysis, CPMsi jointly analyzes all spectra and detected *m/z* signals to identify regions of spectral similarity in a fully data-driven manner. Region-specific molecular features can be extracted from these data-driven regions, enabling general downstream feature analyses and tissue comparisons within CPMsi, as well as export of the results for external analyses.

**Figure 1.**
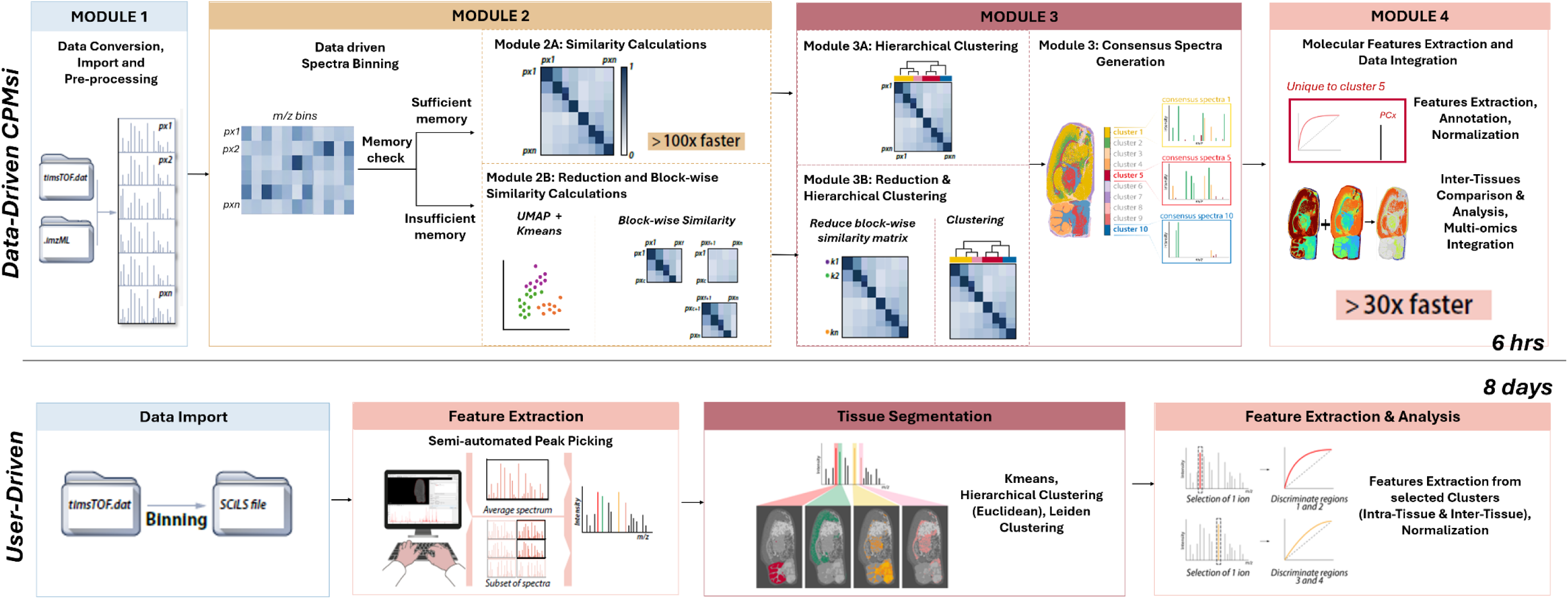
Overview of the CPMsi data-driven workflow and comparison with a conventional user-driven MSI analysis workflow. The upper panel illustrates the CPMsi data-driven workflow. Module 1 performs data conversion, import, and preprocessing (i.e. intensity scaling and filtering) of MSI datasets in parallel. Module 2 applies data-driven spectral binning and automatically evaluates available memory resources. Depending on computational requirements, CPMsi either performs similarity calculations of the full dataset (Modules 2A and 3A) or activates a low-memory strategy (Modules 2B ad 3B) that performs UMAP the binned intensity matrix followed by k-means clustering, and block-wise similarity matrix computation of the full binned intensity matrix. Module 3 performs hierarchical clustering and generates consensus spectra for each cluster. Module 4 enables molecular feature extraction, annotation, normalization, multi-omics data integration, and inter-tissue comparative analyses. The lower panel depicts a conventional user-driven MSI workflow, where data are imported into dedicated software, followed by semi-automated peak selection, tissue segmentation based on the reduced list of peaks, and feature extraction from manually selected regions or clusters. Compared with the user-driven approach, CPMsi minimizes manual intervention while providing a scalable framework (30 times faster) for automated tissue segmentation, consensus spectrum generation, feature extraction, and multimodal data integration across large and complex MSI datasets.

### Computational Workflow

#### Module 1 – Data Conversion, Import and Pre-processing

This module includes three main functionalities: (1) conversion of raw imaging data into the *imzML* format, (2) import of *imzML* datasets and extraction of spectral and spatial information, and (3) spectral preprocessing, including intensity scaling, noise-reduction filtering, and peaks removal. TIMSCONVERT ^16^ is used to convert timsTOF MSI datasets into *imzML* files, but data from other instruments can be imported following external conversion to the *imzML* format. *pyimzML* and *matchms* ^17^ are used for data import and spectral preprocessing. This module also includes a custom peak-removal function which enables the selective exclusion of peaks such as internal standards, matrix-related signals, or other known contaminants.

#### Module 2 – Spectra Binning and Similarity

Here we generate the spectrum-to-spectrum similarity matrix using two alternative computational workflows (2A or 2B), depending on the available system memory. Preprocessed spectra are first aligned into a common *m/z* feature space using a custom data-driven binning strategy. This strategy was designed to correct for small *m/z* variations across spectra and accelerate pixel-to-pixel similarity calculations by transforming spectral data into a matrix representation suitable for efficient vectorized operations. Unlike conventional binning methods, which rely on fixed user-defined *m/z* bin windows, our approach automatically determines the optimal *m/z* reference axis and bin width directly from the data. Further details are provided in the Methods section.

Following binning, spectrum-to-spectrum similarities are computed through fast matrix multiplication of the normalized binned intensity matrix and its transpose, generating a cosine similarity matrix (Equation 1). For datasets that fit into memory, this operation is performed all at once in Module 2A, and the resulting cosine matrix transferred to Module 3A. For datasets exceeding available memory, Module 2B breaks down the calculation and generates three outputs: i) a block-wise cosine similarity matrix, ii) a low-dimensional representation of the binned intensity matrix using UMAP, and iii) k-means clustering of the resulting UMAP embeddings. These outputs are subsequently transferred to Module 3B for scalable hierarchical clustering. Validation of these workflows is provided in Supplementary Information (SI1), including comparison of CPMsi-derived cosine similarities with those obtained using *matchms* to verify the accuracy of our binning strategy. CPMsi showed excellent agreement with the *matchms* implementation while achieving an 118-fold increase in computational speed. Furthermore, CPMsi more effectively preserved global spectral relationships (Supplementary Figure S1).

#### Module 3 - Spectra Clustering and Consensus Spectra Generation

This module provides tools for i) hierarchical clustering, ii) tissue cluster visualization, and iii) consensus spectra generation. Depending on whether the fit-in-memory (Module 2A) or out-of-memory (Module 2B) workflow was used to compute spectral similarities, Module 3 automatically applies the corresponding clustering strategy. For datasets processed through Module 2A, the full cosine similarity matrix is used to identify groups of molecularly similar pixels by standard hierarchical clustering (Module 3A). For datasets processed through Module 2B, a scalable hierarchical clustering approach has been implemented to preserve global spectral relationships in a reduced matrix representation (Module 3B) (example in Figure 2). In both workflows, clustering results are visualized by mapping the cluster label to the spatial coordinates of each spectrum.

**Figure 2.**
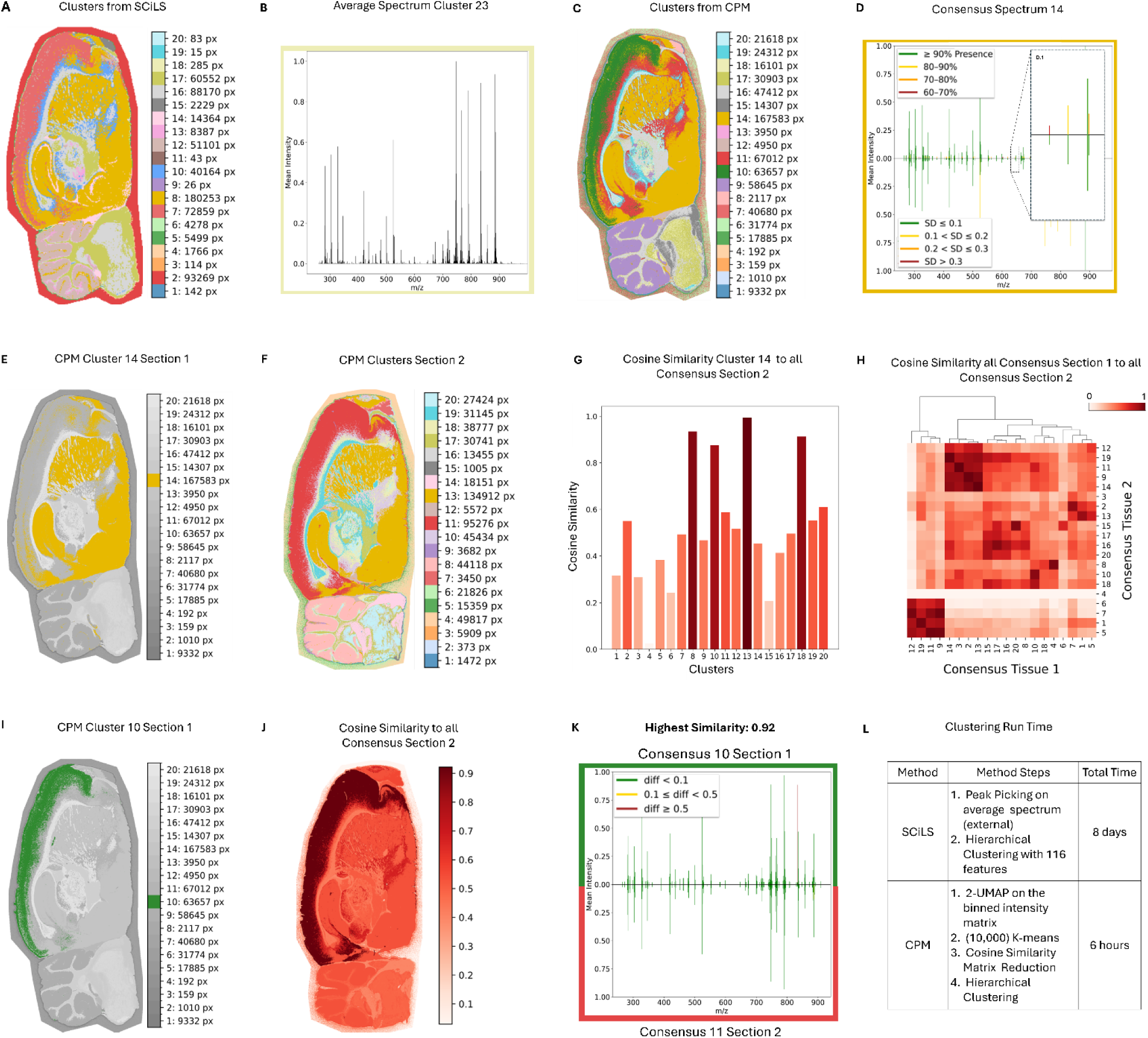
CPMsi on the full dataset. **(A)** Hierarchical Clustering from SCiLS **(B)** Average Spectrum Cluster 23 SCiLS **(C)** Hierarchical Clustering from CPMsi **(D)** Consensus Spectrum CPMsi Cluster 14. Peaks in the top part are colored by their presence across all spectra of the cluster: from green if present more than 90 %, to red when presence is between 70 and 60 %; in the bottom the same peaks are colored by their standard deviation (sd): from green if less than 0.1, to red if above 0.3. The final consensus spectrum will include only the green peaks from both presence and sd. (D.1) zoom in to highlight the different peaks scores. **(E)** CPMsi Cluster 14 Section 1 **(F)** CPMsi Hierarchical Clustering on consecutive section 2 **(G)** Cosine Similarity Matrix between 20 Consensus Spectra from consecutive tissue sections. This allows to map consensus spectra from different datasets. **(H)** Similarity of Consensus Spectrum Cluster 10 CPMsi Section 1, to all other Consensus Spectra from the CPMsi applied to consecutive Section 2. **(I)** Consensus Spectrum Cluster 10 CPMsi Section 1 **(J)** Similarity of Consensus Spectrum Cluster 10 CPMsi Section 1, to all other Consensus Spectra of consecutive Section 2. The highest similarity (0.92) is with the Consensus Spectrum of Cluster 11. **(K)** Mapped Consensus spectrum 10 from section 1 (top) and Consensus Spectrum 11 from section 2 (bottom). Peaks are colored according to their intensity difference: green if the difference is less than 0.1, yellow if it’s between 0.1 and 0.5, red if bigger than 0.5. **(L)** Consensus Spectrum of Cluster 13 Section 2. This Consensus shows the highest cosine similarity with the consensus spectrum of Cluster 14 from Section 1

After clustering, consensus spectra are generated by condensing the spectra within each group. Peaks are retained according to prevalence and intensity stability within the region. This enables region-level representation of molecular profiles with reduction of molecular redundancy. The resulting consensus spectra are used in downstream analyses to increase robustness and computational efficiency.

#### Module 4 – Molecular Features Extraction and Data Integration

Module 4 supports downstream biological interpretation through comparative and integrative analyses of consensus spectra. It also enables the integration of MSI datasets acquired from consecutive tissue sections or multi-omics measurements from the same section (examples in Figure 3 and 4). For inter-tissue comparisons, consensus spectra generated from individual datasets can be merged and treated as a new input for similarity computation and hierarchical clustering (example in Figure 2). This strategy enables the identification of molecular regions that are conserved across tissues or modalities, while highlighting dataset-specific molecular signatures.

**Figure 3.**
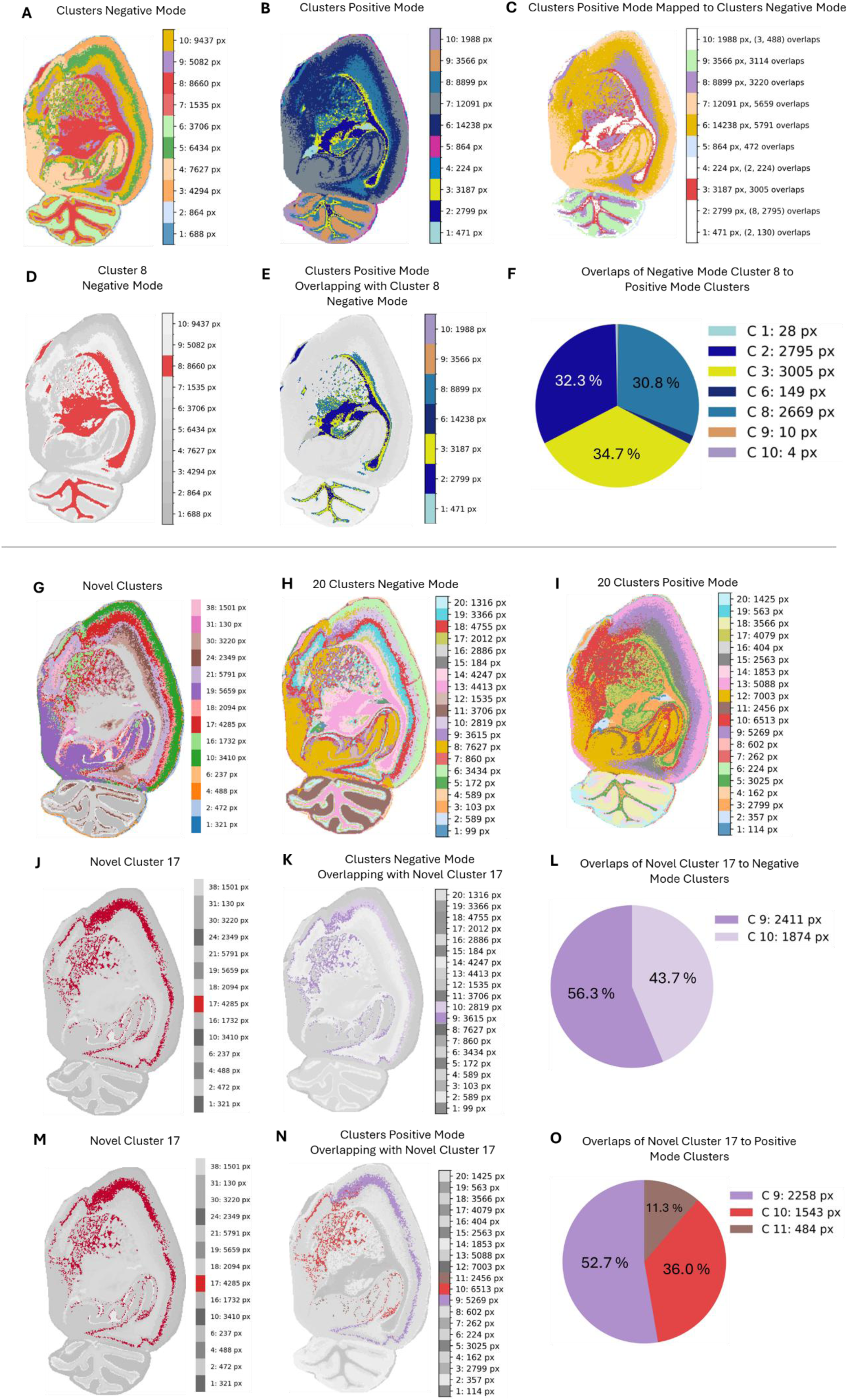
Multi-omics application of the CPMsi. **(A)** Ten clusters of the Negative Mode dataset **(B)** Ten clusters of the Positive Mode dataset **(C)** Spatial overlap between (B) and (A) **(D-E-F)** Example of overlaps: (D) cluster 8 from (A) partially overlaps with clusters 1-2-3-6-8-9-10 from (B) shown in (E). The maximum pixels overlap (34.7%) is with cluster 3 **(G)** Novel Clusters from the combination of 10 Clusters Negative Mode and 10 Clusters Positive Mode. **(H)** 20 clusters of the Negative Mode dataset **(I)** 20 clusters of the Positive Mode dataset **(J)** Novel Cluster 17 **(K)** Overlapping pixels of Clusters Negative Mode with Novel Cluster 17 **(L)** Percentage of overlaps of (K) **(N)** Overlapping pixels of Clusters Positive Mode with Novel Cluster 17 **(O)** Percentage of overlaps of (H)

**Figure 4.**
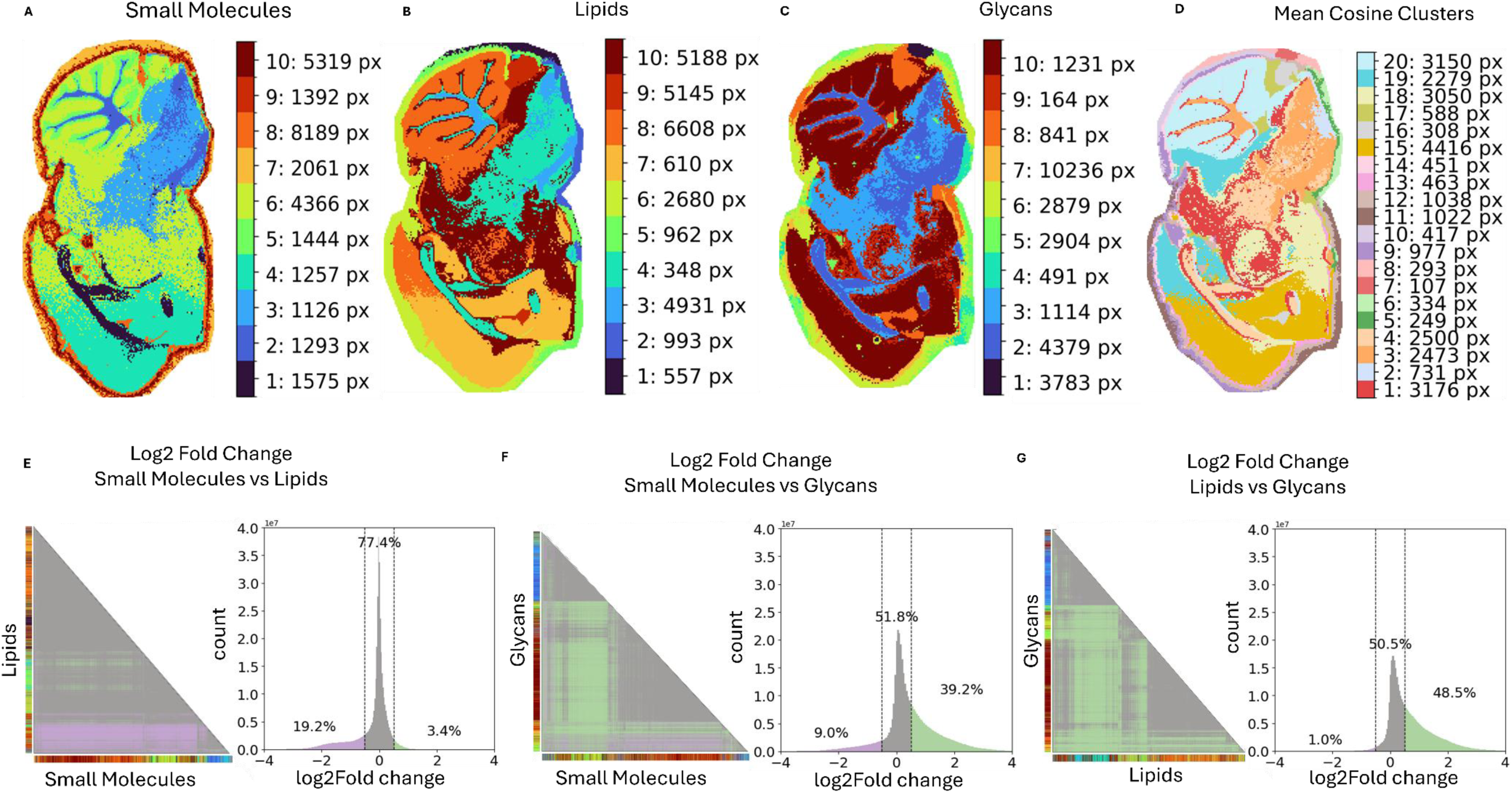
Data Integration using CPMsi. **(A)** Ten clusters of the small molecules dataset **(B)** Ten clusters of the lipids dataset **(C)** Ten clusters of the glycans dataset **(D)** Twenty clusters from the average cosine matrix of the three integrated datasets **(E)** Left: log2 fold change triangular matrix between cosine similarity matrices of small molecules datasets and lipids dataset. Gray area correspond to log2fold change between -0.5 and 0.5; pink area are pixels with values less than -0.5; and green area are above 0.5. Color bars along the sides of the triangle corresponds to the clusters in (A) and (B) respectively; right: histogram of the log2fold change matrix, with percentage values of the number of pixels per value range (left tail values less than -0.5, right tail values more than 0.5) **(F)** Left: log2 fold change triangular matrix between cosine similarity matrices of small molecules datasets and glycans dataset. Color bars along the sides of the triangle corresponds to the clusters in (A) and (C) respectively; right: histogram of the log2fold change matrix, with percentage values of the number of pixels per value range (left tail values less than -0.5, right tail values more than 0.5) **(G)** Left: log2 fold change triangular matrix between cosine similarity matrices of lipids datasets and glycans dataset. Color bars along the sides of the triangle correspond to the clusters in (B) and (C) respectively; right: histogram of the log2fold change matrix, with percentage values of the number of pixels per value range (left tail values less than -0.5, right tail values more than 0.5).

#### Notebooks

The pipeline includes notebooks that provide direct access to its functions for downstream analysis, enabling customized analyses beyond the automated outputs. For example, consensus spectra can be compared across tissue regions to identify shared and region-specific molecular features. In addition, peak-level statistics can be accessed to prioritize prevalent, stable, or discriminatory molecular features. Candidate discriminatory peaks may subsequently be annotated through library matching or by targeted MS/MS analysis (Supplementary Material SI3).

### Intra- and inter-tissue analysis

CPMsi was applied to two consecutive sagittal mouse brain sections measured at 10 µm spatial resolution. 623,599 spectra were generated for the first section, and 587,898 for the second section, both in negative ion mode over an *m/z* range of 260–1100 covering lipids and metabolites. Internal standards (IS) were sprayed as part of our standard MSI acquisition protocol. Raw data conversion into the *imzML* format required 1,46 h running in parallel on an Intel Xeon w7-3545 processor (2.71 GHz). IS peaks, together with their isotopes and adducts, were excluded from the raw spectra, and the corresponding peak intensities saved separately as .csv files for downstream normalization and inter-tissue comparison analysis, not shown in the present work. Computational performance and clustering accuracy were compared with SCiLS Lab 2026 (Bruker Daltonics, Billerica, MA). The data-driven bin width selection for optimal spectra binning was performed on a random subset of 10,000 pixels with bin widths 0.01, 0.015, 0.02, and 0.025 (smaller steps are possible, however for demonstration we only show these four in Supplementary Figure S3). The value 0.02 was automatically selected as optimal with an R² of 0.989. As the spectrum-to-spectrum similarity matrix would require storing on the order of 600,000 × 600,000/2 floating-point values (for the upper triangle), corresponding to 780 GB of memory for float32 or 1,56 TB for float64, calculations were performed using Module 2B, followed by hierarchical clustering implemented in Module 3B. The complete workflow required 6 h for the 623,599 pixels dataset, including similarity computations and clusters generation. In contrast, the equivalent SCiLS Lab analysis required 8 days (Supplementary Information SI2). This gain in computational efficiency did not compromise biological interpretability, as the resulting clusters recovered meaningful hierarchical tissue structures that were comparable to SCiLS Lab segmentation map and remained consistent across the sequential tissue sections (Figure 2). To further investigate discrepancies between the two approaches, we examined cases in which a single SCiLS Lab cluster was divided into multiple CPMsi clusters. Spectral similarity analysis revealed lower similarity among the CPMsi-derived subclusters, indicating the presence of molecular heterogeneity that was not captured by the SCiLS Lab segmentation (Supplementary Figure S4).

While SCiLS represents clusters using average spectra (Figure 2 B) ^18, 19^, CPMsi generates consensus spectra that retain peak-level statistics describing feature prevalence and variability within each cluster (Figure 2 D). Unlike simple averaging, which can preserve high signals originating from a small subset of pixels and obscure their consistency throughout the cluster, consensus spectra explicitly store both the abundance and reproducibility of each peak. This enables the distinction between region-specific molecular features and sporadic peaks arising from matrix effects, or tissue-processing artifacts. Consequently, consensus spectra provide a more robust representation of cluster composition, and support more reliable comparisons across tissue regions or patient samples.

To further demonstrate the potential of consensus spectra for scalable inter-tissue analyses, we compared the 20 consensus spectra generated from each section using cosine similarity (Figure 2 E). This comparison revealed highly similar molecular profiles of corresponding tissue regions, highlighting the reproducibility of the CPMsi workflow across independent MSI acquisitions (Figure 2 F-K). By reducing each dataset to a small set of consensus spectra, CPMsi enables direct comparison of molecularly similar regions while substantially reducing data complexity. Notably, this mapping is independent of spatial coordinates and can therefore be performed across different tissues and measurements. The implications of this approach for region mapping across different tissues are further discussed in the Discussion section.

### Multi-modal MSI data integration using CPMsi

Integration of multi-modal or multi-omics MSI data is supported in Module 4 through two complementary strategies: strategy 1 where cluster identities derived from different data sources are combined to identify novel regions, and strategy 2 where similarity information from multiple sources is combined to investigate molecular crosstalk (see Methods for details).

Here we show applications on two datasets; the first is consecutive lipidomic measurements of mouse brain at 30 µm spatial resolution using dual polarity modes (multi-modal) on the same tissue section with a Bruker timsTOF flex instrument ^20^. The second is a previously published dataset consisting of sequential spatial analysis of the metabolome, lipidome, and glycome (multi-omics), conducted on a sagittal mouse brain section using a Bruker timsTOF flex instrument at 50 µm spatial resolution ^21^.

#### Integration Strategy 1

Two lipidomic ion modes were each analyzed independently, yielding distinct cluster maps (Figure 3 A–B). Partial overlaps were investigated by mapping clusters from the positive-mode dataset onto the negative-mode dataset (Figure 3 C), and vice versa (Supplementary Figure S7), to identify both shared and modality-specific molecular regions. For example, we observed cluster 8 in the negative mode (Figure 3 D) splits into multiple distinct clusters in the positive mode (Figure 3 E), demonstrating complementarity of spatial information across modalities. Cluster-level integration yielded several candidate spatial regions. Following filtering based on consistent subdivision patterns across modalities and comparison with higher-resolution single-modality clustering results (see Methods), a subset of these regions could be fully explained by one of the individual modalities and was therefore excluded (Supplementary Figures S7-S8). Specifically, multimodal clusters identified at a clustering depth of 10 were evaluated against single-modality clustering at a depth of 20 (Figure 3 H-I). Several integrated regions remained distinct from those observed in either modality alone, indicating molecular patterns that emerged only through data integration (Figure 3 J-O).

Consensus spectra were recalculated for the novel clusters, allowing direct comparison with consensus spectra derived from individual modalities. As peak prevalence and intensity statistics depend on the underlying pixel population, changes in the consensus spectra provide a mean to retrieve the additional molecular information captured through multimodal integration. To further characterize the molecular composition of integrated regions, we selected unique *m/z* peaks and performed targeted MS/MS analysis. For example, the feature at *m/z* 766.56 present in the newly resolved cluster 26, was annotated in LIPID MAPS as PE(18:0_20:4) following targeted MS/MS acquisition (Supplementary Information S3).

#### Integration Strategy 2

This is performed by combining similarity information across omics through averaging cosine similarity scores. This strategy captures relationships between molecular-specific clusters and enables direct comparison of spatially resolved molecular organization across the different omics layers (Figure 4 A–C). We used pairwise log2 fold changes between cosine similarity matrices (Equation 2) to assess how pixel-to-pixel molecular relationships differed between modalities, e.g. cosine similarity matrix metabolomic versus cosine similarity matrix lipidomic (Figure 4 D–F). This provides a measure of relative similarity: values around zero indicate pixel pairs with comparable similarity in both modalities, whereas positive or negative values indicate pixel pairs that are more strongly related in one modality than in the other. Here, log2 fold-change values greater than 0.5 and smaller than -0.5 were considered as differing pixel similarity relationships across different omics measurements. Metabolomics and lipidomics showed highly concordant similarity structures, with 77.9% of pixels having the same similarity relationship across the two modalities. Only 3.4% of pixels displayed positive log2 fold change values (> 0.5), whereas 19.2% showed negative values (< - 0.5) associated with inverted patterns between metabolomics and lipidomics, i.e. pixels that are similar in their metabolome lose their similarity at the lipidome level. By contrast, comparisons with glycomics data showed substantially greater differences, with 51.8% and 48.5% of pixel similarities positively differing from lipidomics and metabolomics, respectively. To further quantify agreement between modalities, we calculated an orthogonality score as the angular distance between centered flattened similarity matrices, normalized by 90° (π/2 radians) (Equation 3). By centering the pairwise similarities, the score captures agreement in relative similarity patterns independent of absolute similarity value. A score of 0 indicates identical patterns, a score of 1 indicates orthogonality, and values greater than 1 indicate opposing patterns. The orthogonality scores were 0.68 (corresponding to 61.2°) for metabolomic versus lipidomic, 0.72 (64.8°) for metabolomic versus glycomic, and 0.63 (56.7°) for lipidomic versus glycomic. These results indicate a moderate amount of common structures despite large changes in similarity magnitude highlighted in the log2 fold-change: lipidomic and glycomic data preserve a similar cluster structure but differ more strongly in the magnitude of pairwise similarities (48.5% of pixels); metabolomic and lipidomic show smaller magnitude changes (19.2%) but less agreement in the ordering of pairwise relationships (i.e cluster structure).

## Discussion

CPMsi is a data-driven computational workflow designed to improve the scalability of MSI data processing and analysis, while minimizing user-induced bias. The pipeline integrates advanced spectral binning, similarity scoring, hierarchical clustering, consensus spectra generation, and visualization within a unified and modular framework.

The modular design supports both automated end-to-end or flexible execution of individual modules for targeted analyses. End-to-end execution includes raw data conversion within the pipeline, differently from existing MSI analysis frameworks like Cardinal ^12^, M2aia ^13^, uMaia ^14^, and MSIFlow ^15^, which rely on externally converted *imzML* files. With this functionality, CPMsi reduces user intervention, and supports parallel processing of multiple samples and therefore, scalable analysis of large MSI cohorts. Beyond advantages in workflow execution, the modularity also promotes maintainability and extensibility, facilitating the incorporation of new functionalities, and adaptation to evolving analytical needs. For example, although not explored in this work, future implementations already under development include quantitative analysis tools leveraging internal standards, and automated molecular annotation workflows.

Scalability is further enhanced by the CPMsi’s spectral binning strategy, which effectively aligns spectral signals across all spectra, correcting small mass shifts arising from instrument instability and tissue-height variations, without the need for computationally intensive alignment algorithms ^9, 22^. By transforming the data into a common feature space, this approach also enables highly efficient matrix-based similarity calculations. As demonstrated in our correlation analyses, similarity scores derived from the binned representation showed excellent agreement with established peak-based methods while reducing computational time by more than 100-fold. Clustering analyses further confirmed that the optimal binning width preserves spatially coherent tissue structures. Importantly, these performance gains are achieved without sacrificing spectral information, as CPMsi uses all acquired spectra with the complete set of detected *m/z* features when constructing the similarity matrix. Since for high-spatial-resolution MSI datasets, such as those analyzed in this work (500,000+ pixels), construction of the full pixel-to-pixel similarity matrix becomes impractical due to memory constraints, CPMsi includes an alternative block-wise similarity workflow (Module 2B and 3B). This workflow partitions computations into manageable subsets and stores intermediate results on disk. Combined with a hierarchical reduction strategy, it enables clustering of large MSI datasets while preserving global spectral relationships. Although this approach incurs additional overhead and is therefore slower than in-memory processing (Supplementary S1), the successful analysis of our dataset demonstrates the ability of CPMsi to extend similarity-based MSI analysis beyond memory limitations without information loss.

The effectiveness of this full spectrum-to-spectrum similarity representation is further enhanced by the use of cosine similarity, which distinguishes CPMsi from workflows based on Euclidean distance, such as the hierarchical clustering implemented in SCiLS Lab. Euclidean distance can perform well on reduced peak-picked datasets commonly used in MSI analysis; however, dimensionality reduction and feature selection inevitably simplify molecular complexity and may obscure subtle heterogeneity within tissue regions. This effect was evident in our analyses, where regions represented as a single cluster by SCiLS were resolved into multiple molecularly distinct subclusters by CPMsi. Additionally, cosine similarity compares spectra based on their orientation in feature space rather than their absolute distance, making it particularly well suited for identifying pixels with similar molecular compositions in high-dimensional MSI data ^23^. Despite these advantages, cosine-based spectrum-to-spectrum analysis is rarely performed at the pixel level because of its substantial computational demands. CPMsi overcomes this limitation through efficient spectra binning and matrix-based computation, enabling cosine similarity analysis across hundreds of thousands of spectra and combining improved molecular resolution with the scalability required for large-scale MSI studies.

The similarity-based framework also provides the basis for the generation of consensus spectra from molecularly related tissue regions. Unlike traditional MSI analyses that rely on average spectra ^4, 15, 24^, CPMsi provides a more robust characterization of regional molecular signatures through consensus spectra, which capture peak occurrence frequencies and intensity variability. In the present study, we showed that consensus spectra enable cross-sample comparisons by mapping molecularly similar regions across tissue sections with different spatial coordinates. Because this representation is based on molecular similarity rather than tissue geometry, it can be readily extended to cohort-level analyses, allowing corresponding regions to be identified across samples from different patients. Such capabilities will ultimately support the development of MSI tissue atlases and disease-specific reference databases, requiring acquisition and analysis of large number of datasets.

CPMsi uses the similarity-based framework also for integration of multi-modal or multi-omics MSI datasets, enabling the identification of tissue heterogeneity that may remain unresolved within individual molecular layers. Our analysis on the multi-modal lipidomic dataset confirmed that each modality captures distinct aspects of the lipidomic landscape of the mouse brain ^25^. In particular, the observed splitting of clusters across modalities reflects molecular heterogeneity within tissue compartments that appear homogeneous when examined using a single acquisition mode. Importantly, integrated analysis revealed regions that were not resolved in either modality alone, highlighting the value of combining complementary molecular information. A similar integration strategy was recently shown by SAMI ^20^; however, rather than relying primarily on low-dimensional embeddings for multimodal clustering, CPMsi combines full-spectrum cosine similarity matrices across modalities, preserving molecular relationships between pixels and enabling tissue regions to emerge directly from the shared similarity structure. In addition, CPMsi supports cluster-level integration which combines cluster assignments from individual modalities to define integrated tissue regions.

Together, these results demonstrate that CPMsi enables efficient and scalable analysis of large MSI datasets while preserving the full molecular complexity of acquired spectra. By supporting standardized tissue characterization, cross-sample comparisons, and multi-modal/multi-omics data integration, CPMsi provides a framework for the systematic analysis of MSI cohorts. These capabilities will facilitate biomarker discovery and support the development of robust molecular classification models with potential diagnostic and prognostic applications in clinical routine. Importantly, MSI provides a complementary view of tissue organization by enabling the direct investigation of molecular interactions rather than relying on interactions inferred from gene expression patterns.

## Materials and Methods

### Chemicals

N-(1-Naphthyl)ethylenediamine dihydrochloride (NEDC), methanol (MeOH) ultra liquid chromatography–mass spectrometry (LC–MS) (Actu-all chemicals, The Netherlands), acetonitrile (ACN) ultra LC–MS (Actu-all chemicals, The Netherlands), Milli-Q water (MQ) (EMD Millipore Corp), MSI splash mix (Avanti, USA).

### Sample collection

Brains were collected from control 6–8-week-old OF1 male mice (Charles River Laboratories) and snap-frozen in liquid nitrogen under the AVD project license number 11600202216547.

The multi-modal (negative and positive ion mode) lipidomic datasets were previously acquired as described in ^26^. The multi-omics datasets were downloaded as *imzML* files via the public repository in ^21^.

### Sample preparation for MALDI MSI of mouse brains

Sagittal mouse brain sections of 10 µm thickness were sectioned on a cryostat (CM3050 S, Leica Biosystems) and mounted on ITO slides (Visiontek Systems Ltd). The ITO slide was then vacuum freeze-dried for 15 min. First, 20 layers of MSI splash mix (Avanti, USA) in absolute methanol was sprayed on the tissues using the HTX M3+ sprayer (HTX Technologies, LLC) with the following parameters: 60°C, 80 µL/min flow rate, 1200 mm/min velocity, 2 mm track spacing, CC pattern and 10 s drying time between each layer. Subsequently, 20 layers of 7 mg/mL N-(1-Naphthyl)ethylenediamine dihydrochloride (NEDC) matrix in 70% MeOH/25% ACN/5% MQ was sprayed on the tissues using the HTX M3+ sprayer (HTX Technologies, LLC) with the following parameters: 60°C, 80 µL/min flow rate, 2000 mm/min velocity, 3 mm track spacing, CC pattern and 30 s drying time between each layer.

### MALDI MSI

MALDI-TOF-MSI measurements were performed on a MALDI-2-timsTOF flex instrument. Negative ion mode spectra with a mass range of *m/z* 260-1100 were acquired with 6 µm beam scan resulting in a final spatial resolution of 10 µm. High spatial resolution images were obtained with 64% laser power and 100 shots per pixel at a speed of 10 kHz in MALDI and 1 kHz in MALDI-2 mode (trigger delay 5.0 µs). Prior to data acquisition the instrument was calibrated using red phosphorus. Data acquisition was performed using timsControl and flexImaging 7.3.

### Computational Hardware

All computations were performed on Windows workstation (HP z8 Fury) equipped with an Intel Xeon w7-3545 processor, 512 GB RAM, and an NVIDIA RTX A4000 GPU with 16 GB VRAM.

### Software implementation

The CPMsi is built on top of various Python libraries with custom algorithms to enable scalable analysis of large-scale MSI data. High computational efficiency is achieved through vectorized operations and parallelization across all modules. The framework adopts a modular architecture composed of independent processing modules that can be executed either individually or sequentially as a complete workflow. Each module has a standardized structure that is compatible with MZmine ^27^:

- Functions: reusable source code implementing core computational procedures;
- Tasks: workflow-specific execution logic;
- RunTasks: command-line interfaces for module execution;
- Parameters: YAML configuration files containing user-defined settings and processing inputs;

Automated execution of the complete workflow is controlled through the *`run_pipeline.py`* using user-provided YAML configuration file. This design promotes reproducibility by separating configuration settings from processing logic.

### Module 1: Data Conversion, Import and Pre-processing

#### Module 1.1: Data Conversion

This step is required when raw vendor-specific data need to be converted into the *imzML* standard format. We use TIMSCONVERT ^16^ to convert timsTOF MSI datasets into *imzML* files. This process is embarrassingly parallelized through a custom Python script that maximizes the utilization of available CPU cores, thereby reducing processing times and improving scalability. This facilitates efficient processing of large cohort studies.

#### Module 1.2: Data Import

*pyimzML* is used to import *imzML* files. Spectral data are extracted together with their spatial coordinates using `getspectrum()`; each spectrum is converted into a *matchms* Spectrum object containing the m/z values, intensity values, and a unique spectrum identifier. The *x* and *y* coordinates are stored separately in a dictionary object.

#### Module 1.3: Spectral Pre-processing

*matchms* filtering module is used to perform optimized spectra intensity scaling (`filtering.normalize_intensity()`), and noise-reduction filtering with a user-defined intensity threshold (`filtering.select_by_relative_intensity()`). Peak removal is performed using `remove_peaks_from_list()`. This function implements a nearest-neighbor search algorithm to identify and remove undesired peaks, such as those corresponding to internal standards. The algorithm also supports the removal of adduct and isotopic species through correlation analysis. Peaks whose spatial distributions exhibit a correlation with the parent ion above user-defined thresholds are automatically identified as related species and removed from the spectra. Separate correlation thresholds for adducts (`adduct_corr_threshold`) and isotopes (`isotope_corr_threshold`) are provided as input parameters to the function, with default values of 0.80 and 0.90, respectively. The intensity values of all removed peaks are stored separately in a *csv* file, allowing their subsequent retrieval for downstream applications such as peak normalization and quantification in case the removed peaks are internal standards.

### Module 2: Spectra Binning and Similarity

Module 2 generates the binned intensity matrix and the spectrum-to-spectrum similarity matrix. It includes two execution branches depending on the size of the dataset and available computational resources, or on user choice: a standard fit-in-memory workflow (Module 2A) and an out-of-memory workflow (Module 2B). Both branches use the same principles for spectra binning but differ in how the similarity matrix is computed and stored. In addition, Module 2B performs UMAP on the binned intensity matrix and k-means.

#### Module 2.1: Data-Driven Spectra Binning

Spectral binning was performed to match peaks across spectra and facilitate spectrum-to-spectrum similarity calculations. Conventional binning methods divide the *m/z* range into fixed intervals and assign peak intensities to a predefined reference grid. This approach may result in peak splitting when signals fall near bin boundaries, and it requires users to define a fixed *m/z* axis. To address these limitations, a data-driven binning strategy was implemented in the `create_intensity_matrix()` function. This function performs a two-steps binning and optimizes the selection of representative *m/z* bins.

##### 2.1.1 Binning Step 1

In the first step, all spectra are conventionally binned onto a uniform *m/z* axis spanning the global minimum and maximum *m/z* values observed in the dataset. The spacing between consecutive bins is determined by the `bin_width` parameter, expressed in *m/z* unit. For each spectrum, peak intensities associated with the same bin are aggregated to generate a binned intensity representation.

To reduce the influence of poorly supported signals and noise, bins detected in fewer than 10% of spectra are excluded.

##### 2.1.2 Binning Step 2

In the second step, the binning procedure is repeated with an offset in the *m/z* range of half bin width. For example, when using a bin width of 0.01 *m/z*, the second grid is generated with an offset of +0.005 *m/z*. This shifted binning captures peaks that lie near bin boundaries in the first grid and may therefore be split across adjacent bins.

##### 2.1.3 Selection of Representative Bins

The two binned intensity matrices are subsequently compared using a custom sliding-window algorithm `(_select_bins_local_winners_sliding_window()`). This algorithm merges the two matrices, evaluates consecutive bins within local windows of `window_width`, and selects the ones with the highest spectral coverage. This logic favors bin configurations that minimize peak splitting, as signals represented by a single bin generally exhibit higher coverage than those distributed across adjacent bins. To avoid retention of bins representing the same molecular feature, neighboring bins with equal spectral coverage are further compared using the Pearson correlation of their intensity profiles. Bins with correlation exceeding a user-defined threshold are considered redundant, and only the bin with the higher total intensity is retained.

Importantly, the `bin_width` and the sliding `window_width` parameters are independent. Whereas the `bin_width` determines the resolution of the initial binning process, the `window_width` defines the local neighborhood examined during the selection of representative bins and therefore affects the detection of isotopic peaks and adducts.

##### 2.1.4 Final Binning

The final set of representative bins selected through the sliding-window procedure constitutes a non-uniform, *m/z* reference grid optimized for the dataset. Consequently, all spectra are re-binned onto this final reference using a *Numba*-accelerated binary-search algorithm. The resulting intensity matrix is used for spectral similarity calculations and downstream analyses.

#### Module 2.2: Spectra Similarity

Following binning, spectrum-to-spectrum similarity calculations are performed through efficient matrix multiplication using either Module 2A or Module 2B. This calculation produces a symmetric similarity matrix of size *n×n*, where *n* is the number of spectra. Because memory requirements of the similarity matrix increase quadratically with *n*, its expected size is estimated prior to computation, and the appropriate execution strategy is selected automatically. For example, a dataset of 500,000 spectra would generate a similarity matrix comprising 250 billion pairwise similarity values (500,000 x 500,000). Even stored as 32-bit floating-point values, this matrix requires approximately 1 TB of memory, far exceeding the capacity of a typical workstation equipped with 64 GB RAM. Such dataset would therefore be processed using the out-of-memory workflow (Module 2B).

##### 2.2.2A In-Memory Similarity Calculation

Standard pairwise spectra similarity is computed in Module 2A using a custom function implemented in JAX ^28^. The algorithm simultaneously calculates cosine similarity score for all pairs of spectra through matrix multiplication:

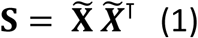

where **S** ∈ ℝ^*n* ×*n*^ is the similarity matrix of size *n* spectra × *n* spectra, **X̃** is the normalized intensity matrix **X** ∈ ℝ^*n* ×*p*^ of size *n* spectra × *p* bins, and ***X̃***^T^ is the transpose. Each entry **S**_*ik*_ of **S**, represents the cosine similarity score between normalized spectra **x̃**_*i*_ and **x̃**_*k*_ of **X̃**, with

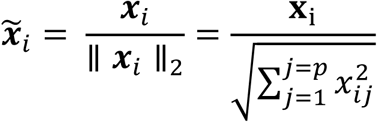

Equation 1 can also be performed with GPU acceleration.

##### 2.2.2B Out-of-Memory Similarity Calculation

Spectra similarity is computed in blocks for larger datasets that do not fit into memory. In this implementation, the normalized intensity matrix is first divided into two equal partitions, and the similarity calculation is done in three blocks: two squares within partition blocks and one rectangular between partitions block (Supplementary Figure S10). Each block is computed using tiled matrix multiplication on a GPU. Row and column subsets are transferred to the GPU in fixed-size tiles, multiplied to obtain local similarity matrices, and written into their corresponding locations within the output block. For square blocks, only upper-triangular tiles were explicitly evaluated, and the lower-triangular region was reconstructed by symmetry. Each completed similarity block is stored independently as a compressed *NumPy* array for downstream analysis. The procedure is summarized in Supplementary Information (Algorithm 1).

Module 2B includes additional steps necessary to circumvent the computational limitations associated with large cosine similarity matrices and enable hierarchical clustering in Module 3B. These steps consist of:

1. Dimensionality reduction on the intensity matrix: UMAP function implemented in the *umap-learn* Python library is used to project the binned intensity matrix into a 2- or 4-dimensional space.
2. Clustering in the embedded space: Mini-Batch K-means ^29^ is applied on the low-dimensional representation to obtain *k* coarse similar groups. Cluster assignments are stored in a dictionary object for later use in cosine matrix compression. The value of *k* is automatically calculated from the target reduction level specified by the user in the YAML configuration file (e.g. 50-fold reduction).

#### Module 2.3: Data-Driven Selection of bin_width

To minimize user-defined parameters and enable a fully data-driven workflow, the optimal `bin_width` for binning is determined automatically. The binning and similarity calculations described above are iteratively applied across a range of candidate values (e.g., 0.001-0.03 in increments of 0.005), using a random subset of spectra. For each candidate bin width, similarities derived from the resulting binned intensity matrix are compared with reference similarities obtained using the `CudaCosineGreedy()` function from *matchms* ^17, 30^. This GPU-accelerated method efficiently approximates peak matching while closely reproducing cosine similarity scores from the exact *matchms* implementation. Pearson correlation between the reference and CPMsi similarity scores is then calculated, and the bin width value yielding the highest correlation is selected as optimal and subsequently used for the analysis.

### Module 3: Spectra Clustering and Consensus Spectra Generation

Module 3 implements two hierarchical clustering workflows, Module 3A and Module 3B, designed for cosine similarity matrices generated by Modules 2A and 2B, respectively. Consensus spectra are subsequently computed for all identified clusters using a unified workflow. The module further includes tools for visualizing clustering results and consensus spectra.

#### Module 3.1: Hierarchical Clustering

##### 3.1.3A In-memory Hierarchical Clustering

When sufficient memory is available, hierarchical clustering is performed directly on the full cosine similarity matrix using *SciPy.* Cosine distances are calculated as `1–cosine` `similarity`, and a linkage matrix (matrix representation of the hierarchical tree) is computed using the ward method. Cluster assignments are obtained using the `fcluster()` function with `maxclust` criterion, which cuts the hierarchical tree at the height that gives the user-defined number of clusters (e.g., 10).

##### 3.1.3B Out-of-memory Hierarchical Clustering

For datasets where the full similarity matrix cannot be loaded into memory, clustering is performed using a compressed similarity representation derived from the stored similarity blocks.

First, the precomputed k-means cluster assignments are used to reduce the dimensionality of the similarity matrix. For each block file, cosine similarity scores between spectra belonging to the same k-means cluster are aggregated to form a reduced cluster-level similarity matrix. To preserve the geometric interpretation of cosine similarity, aggregation is performed in angular space rather than directly on cosine values. Specifically, cosine similarities are transformed into angles, averaged, and then converted back to cosine space:

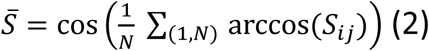

where *S*_*ij*_ denotes the cosine similarity between spectra *i* and *j*, and *N* is the number of similarity values being averaged.

The resulting compressed similarity matrix is subsequently used as input for hierarchical clustering in *SciPy*. Final cluster labels are propagated to individual spectra through the corresponding k-means cluster assignments.

#### Module 3.2: Consensus Spectra Generation

`define_consensus_spectra()` computes for each cluster, peak prevalence, mean intensity, and intensity variability across spectra. These statistics are stored in the consensus spectrum together with a consistency score reflecting the reproducibility of each peak within the cluster. Peaks can be visualized in the spectrum using a color scale ranging from green (high prevalence and low variability) to red (low prevalence and/or high variability), enabling rapid assessment of feature reliability while preserving the full molecular complexity of the region.

This function can be applied to any selected set of pixels, providing additional functionalities beyond intra-tissue cluster analyses. Consequently, the pipeline can be extended to applications such as inter-tissue comparisons, and investigation of novel spatial regions identified from multi-modal and multi-omics data integration.

#### Module 3.3: Cluster Visualization

Cluster labels are automatically mapped back to their corresponding spectra and spatial coordinates for spatial visualization, enabling visual inspection and direct comparison with complementary imaging data such as histology or anatomical atlases. We provide functions to visualize either the complete clustering solution or selected clusters.

Two visualization schemes are available. First, a discrete color palette assigning warm yellow and orange tones to larger clusters, facilitating visual perception of dominant tissue regions. Second, a Turbo-based continuous palette colors clusters according to the similarity of their consensus spectra, allowing relationships between clusters to be explored directly within the spatial context.

### Module 4: Molecular Features Extraction and Data Integration

Module 4 supports downstream biological interpretation through comparative analysis of consensus spectra and integration of complementary datasets.

#### Module 4.1: Comparative Analysis of Consensus Spectra

Consensus spectra generated in Module 3 can be compared within and across datasets to identify biologically meaningful molecular features.

*Intra-tissue analysis.* Cluster-specific molecular features are identified by investigating consensus spectra similarity and peaks statistics. Peaks uniquely enriched in individual clusters can be extracted and further investigated through spectral library searching and annotation workflows.

*Inter-tissue analysis.* Consensus spectra originating from different tissues, or consecutive sections of the same tissue can be analyzed jointly to identify conserved tissue structures and molecular signatures. In this analysis, consensus spectra from all datasets are combined into a single dataset and treated as new input for the pipeline. The combined dataset is subsequently processed using Modules 2 and 3, enabling direct comparison based on spectral similarity and identification of shared and tissue-specific molecular features.

#### Module 4.2: Multi-modal and Multi-omics Data Integration

Integration functions enable integration of multiple MSI modalities acquired from datasets sharing a common spatial coordinate framework. Two complementary integration strategies are supported.

##### Module 4.2.1 Integration Strategy 1 for the identification of novel spatial regions

This strategy is designed to identify novel spatial regions that emerge only through multimodal integration and are not detected in the individual modalities. First, cluster assignments obtained independently for each modality are combined at matching spatial coordinates to generate integrated cluster labels. For example, if pixel *p*_*i*_belongs to cluster *n* in modality 1 and cluster *m* in modality 2, it is assigned to the integrated cluster *nm*. Each integrated cluster therefore represents a unique combination of cluster identities across modalities. Candidate novel regions are identified using a two-steps procedure.

*Step 1.* For each modality, the redistribution of pixels from the original clusters to the integrated clusters is quantified using a column-normalized contingency table. An original cluster is considered to undergo a structured subdivision when its pixels redistributed between 2 and ***N*** integrated clusters (***N*** = 5 by default), with each integrated cluster receiving at least a minimum fraction ***f*** of the original cluster (***f*** = 0.05 by default). Integrated clusters participating in such subdivisions are identified independently for each modality. Only integrated clusters supported by structured subdivisions in all modalities are retained as candidate novel regions. This criterion excludes both trivial one-to-one correspondences and highly fragmented redistributions.

*Step 2.* Candidate regions are subsequently compared with deeper clustering results obtained independently for each modality. For example, candidates derived from clustering with ***k*** *= 10* are compared against clustering results obtained with ***k*** *= 20*. For each candidate integrated cluster, the proportion of pixels assigned to each deeper cluster is calculated, and the maximum overlap fraction is determined. Candidates are excluded if more than 80% of their pixels overlap with a single deeper cluster in any modality. Regions surviving this filtering step are considered novel spatial domains revealed by multi-modal integration rather than by increased clustering resolution within an individual modality.

A schematic overview of the cluster-level integration strategy is provided in Supplementary Figure 9.

##### Module 4.2.2 Integration Strategy 2 for the investigation of molecular relationships

This strategy integrates molecular information at the similarity-matrix level before clustering. Cosine similarity matrices are computed independently for each modality and transformed to angular space (Equation 2), where similarities are aggregated by averaging. The resulting consensus matrix is then converted back to cosine space and processed using the clustering workflow described in Module 3. The resulting clusters represent spatial domains supported by the combined molecular information from all integrated modalities.

##### Module 4.2.3 Similarity Analysis Tools

*Log2 Fold Change.* To compare the spatial organization captured by different omics modalities, pairwise differences between modalities are quantified using the log2 fold change of corresponding similarity matrix entries:

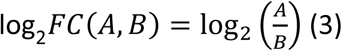

where *A* and *B* denote the similarity matrices, therefore the division is element-wise:

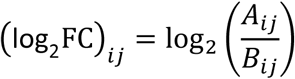

with *A*_*ij*_ and *B*_*ij*_ indicating the matrix entries *ij*.

Log2 fold-change values close to 0 indicate similar pixel-to-pixel relationships between modalities, whereas larger positive or negative values indicate modality-specific similarity patterns.

*Orthogonality Score.* The orthogonality score is calculated as the angular distance between similarity matrices using:

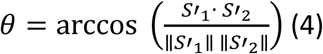

where *S*′_1_and *S*′_2_ are the centered flatten similarity matrices from two datasets:

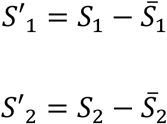

with *S̅*_1_ and *S̅*_2_ denoting the means of S1 and S2, the flatten similarity matrices.

In this representation, the upper triangular elements of each cosine similarity matrix, excluding the diagonal, are centered by subtracting their mean and treated as vectors. An orthogonality score of 0 indicates identical similarity structures, whereas larger angles indicate increasing dissimilarity between modalities.

## Code availability

https://github.com/MassSpectrometryImaging/CPMsi

## Data availability

The data is available from the authors upon request.

## Acknowledgements

This work is supported by a grant from the Dutch Kidney Foundation Kolff+ number 21OK+15. TJR and MG were partially supported by The Novo Nordisk Foundation Center for Stem Cell Medicine (reNEW), supported by Novo Nordisk Foundation grants (NNF21CC0073729). TJR is supported by the European Union through ERC grant (SPARK 101140863). Views and opinions expressed are however those of the author(s) only and do not necessarily reflect those of the European Union or the European Research Council Executive Agency. Neither the European Union nor the granting authority can be held responsible for them. We thank Manon Zuurmond (Department of Internal Medicine, LUMC, Leiden, the Netherlands) for discussion and feedback on Figure 1.

## References

1. Clark, A. et al., Rapid discrimination of pediatric brain tumors by mass spectrometry imaging. J Neurooncol. (2018 Nov).

2. Phillips, L., Gill, A. & Baxter, R., Novel Prognostic Markers in Triple-Negative Breast Cancer Discovered by MALDI-Mass Spectrometry Imaging. Front Oncol. 9 (2019 May 14).

3. Colley ME, Esselman AB, Scott CF, Spraggins JM, High-Specificity Imaging Mass Spectrometry. Annu Rev Anal Chem (Palo Alto Calif) 17 (1), 1–24 (2024 July).

4. Nunes, J. et al., Integration of mass cytometry and mass spectrometry imaging for spatially resolved single-cell metabolic profiling. Nat Methods 21 (10), 1796–1800 (2024 Oct).

5. Godfrey, T. et al., Integrating Ambient Ionization Mass Spectrometry Imaging and Spatial Transcriptomics on the Same Cancer Tissues to Identify RNA-Metabolite Correlations. Angew Chem Int Ed Engl 64 (24), e202502028 (2025 Jun 10).

6. Tasca, P. et al., Application of spatial-omics to the classification of kidney biopsy samples in transplantation. Nat Rev Nephrol 20 (11), 755–766 (Nov 2024).

7. Balluff, B., Hopf, C., Porta Siegel, T., Grabsch, H. & Heeren, R., Batch Effects in MALDI Mass Spectrometry Imaging. J Am Soc Mass Spectrom 32 (3), 628–635 (2021 Mar 3).

8. La Rocca, R., et al., Adaptive Pixel Mass Recalibration for Mass Spectrometry Imaging Based on Locally Endogenous Biological Signals. Anal Chem 93 (8), 4066–4074 (2021 Mar 2).

9. Eriksson, J., et al., MSIWarp: A General Approach to Mass Alignment in Mass Spectrometry Imaging. Anal Chem 92 (24), 16138–16148 (2020 Dec 15).

10. Lieb, F., Boskamp, T. & Stark, H., Peak detection for MALDI mass spectrometry imaging data using sparse frame multipliers. J Proteomics 225 (2020 Aug 15).

11. Veselkov, K., et al., BASIS: High-performance bioinformatics platform for processing of large-scale mass spectrometry imaging data in chemically augmented histology. Sci Rep 8 (1) (2018 Mar 6).

12. Bemis, K. A. . F. M. C. . G. D. e. a., Cardinal v.3: a versatile open-source software for mass spectrometry imaging analysis. Nat Methods (20), 1883–1886 (2023).

13. Cordes, J., et al., M2aia-Interactive, fast, and memory-efficient analysis of 2D and 3D multi-modal mass spectrometry imaging data. Gigascience 10 (7) (2021 Jul 20).

14. Schede, H., et al., Unified mass imaging maps the lipidome of vertebrate development. Nat Methods 22 (9), 1981–1994 (2025 Sep).

15. Spangenberg, P., et al., msiFlow: automated workflows for reproducible and scalable multimodal mass spectrometry imaging and microscopy data analysis. Nat Commun 16 (1) (2025 Jan 27).

16. Luu, G. et al., TIMSCONVERT: a workflow to convert trapped ion mobility data to open data formats. Bioinformatics 38 (16), 4046–4047 (2022 Aug 10).

17. Huber, F., Verhoeven, S., Meijer, C. & al., e., matchms - processing and similarity evaluation of mass spectrometry data. J Open Source Softw 5 (52) (2020).

18. Berlin, K., Huizar, C., Garza, C., Forsthuber, T. & Bach, S., MALDI-TOF mass spectrometry imaging of sulfatide lipid expression in the CNS of mice with experimental autoimmune encephalomyelitis. Sci Rep 16 (1) (2026 Feb 28).

19. Dilillo, M., et al., Ultra-High Mass Resolution MALDI Imaging Mass Spectrometry of Proteins and Metabolites in a Mouse Model of Glioblastoma. Sci Rep 7 (1) (2017 Apr 4).

20. Zhan, L., Huang, Y. & Wang, G., Multi-modal mass spectrometry imaging of a single tissue section. J Mass Spectrom 59 (8) (2024 Aug).

21. Clarke, H., Ma, X., Shedlock, C. & al, e., Spatial mapping of the brain metabolome lipidome and glycome. Nat Commun 16 (1) (2025).

22. Johnson, W., Li, C. & A, R., Adjusting batch effects in microarray expression data using empirical Bayes methods. Biostatistics 8 (1), 118–27 (2007 Jan).

23. Smets T, V. N. C. M. et al., Evaluation of Distance Metrics and Spatial Autocorrelation in Uniform Manifold Approximation and Projection Applied to Mass Spectrometry Imaging Data. Anal Chem 91 (9), 5706–5714 (2019 May 7).

24. Alexandrov, T., MALDI imaging mass spectrometry: statistical data analysis and current computational challenges. BMC Bioinformatics (2012).

25. Ellis, S. et al., More from less: high-throughput dual polarity lipid imaging of biological tissues. Analyst 141 (12), 3832–41 (2016 Jun 21).

26. van der Vloet, L. et al., Mass Spectrometry Imaging Reveals the Distribution of a GABRG2 Targeting Antisense Oligonucleotide and Its Functional Effect in Rat Brain. ACS Chem Neurosci 16 (20), 4064–4075 (2025 Oct 15).

27. Schmid, R., Heuckeroth, S., Korf, A. & al, e., Integrative analysis of multimodal mass spectrometry data in MZmine 3. Nat Biotechnol 41 (4447-449) (2023).

28. Zhang, J., JAX: composable transformations of ython+

29. Sculley, D., Web-scale k-means clustering, presented at Proceedings of the 19^th^ international conference on World wibe web, pages 1177-1178, 2010 (unpublished).

30. Onoprishvili, T., Yuan, J., Petrov, K. & al., e., SimMS: a GPU-accelerated cosine similarity implementation for tandem mass spectrometry. Bioinformatics 41 (3) (2025).

31. Tobias, F. & Hummon, A., Considerations for MALDI-Based Quantitative Mass Spectrometry Imaging Studies. J Proteome Res 19 (9), 3620–3630 (2020 Sep 4).

