## supporting information for "CPMsi: a data-driven Computational Pipeline for fast, large-scale Mass Spectrometry Imaging analysis"

### SI1. Performance of CPMsi

To validate our approach, we initially applied CPMsi on a representative subregion of 44,419 pixels before applying the workflow to the full MSI dataset (Figure S1 A). As part of the CPMsi data-driven binning strategy, cosine similarity scores derived from binned spectra are compared with similarity scores calculated using the *matchms* [1] implementation of `CudaCosineGreedy` [2]. This comparison is used to identify the optimal binning width that yields maximum agreement between binned and peak-based similarity calculations. Because strong agreement indicates that the binning procedure faithfully preserves the spectral relationships captured by established peak-based methods, this comparison also serves as an independent validation of the CPMsi similarity framework. At the optimal binning width, correlation analysis demonstrated excellent agreement between the two approaches ($R^2 = 0.972$; Figure S1 B). Under these conditions, CPMsi achieved an 118-fold increase in computational speed relative to the reference *matchms* computation. Agreement in cosine similarity scores is further reflected in the clustering, which showed high pixel-level overlaps (Figure S1 C-E).

In Figure S1 C-D, we highlight a small area where the two methods clearly diverge. To investigate this disagreement, we expanded the subregion to include a broader spatial context comprising 75,428 pixels. In this expanded view, the reference method which previously separated these areas, now grouped them into the same cluster. This shows that the CPMsi more effectively captures global spatial structures, whereas the reference approach is more sensitive to local variations.

To validate the out-of-memory workflows (Modules 2B and 3B), we applied them to the 44,419 pixels region and compared with the results obtained using the standard in-memory workflows (Modules 2A and 3A). Pearson correlation of the two generated cosine similarity matrices demonstrated perfect agreement between the two methods ($R^2 = 1.0$; Figure S2 A). Furthermore, hierarchical clustering maps produced by the standard in-memory and out-of-memory workflows showed highly similar spatial cluster distributions and overall tissue architecture (Figure S2 B-C, Table S1), indicating that the out-of-memory implementation faithfully reproduces the results of the standard in-memory workflow, however it took 92.3 secs against 3.4 secs of the in-memory workflow.

**SI2. Comparison of CPMsi clusters to SCiLS segmentation map**

The lipidomic brain dataset, comprising more than 600,000 pixels, was imported into *SCiLS Lab*. Feature finding was performed using the `T-ReX²(QTOF)` algorithm with the following settings: spectrum mode, *all spectra*; spatial noise filtering, *weak*; spectrum coverage, *5%* (31,179 pixels); maximum number of features, *unlimited*; intensity threshold, *0.1%*; and interval width, *0.02 Da*. This initial feature-finding step yielded 11,491 m/z features.

Hierarchical clustering-based segmentation was subsequently attempted on all individual spectra using this list. However, the computational demands exceeded the capabilities of the system, preventing completion of the analysis. To reduce data dimensionality, the average spectrum was exported and manually curated in *mMass* [3], resulting in the selection of 115 *m/z* features. These features were imported back into *SCiLS Lab*, and segmentation was repeated using the reduced feature set. The clustering analysis completed after approximately 8 days of computation. The resulting hierarchical tree was manually explored, and cluster labels corresponding to a 20-cluster solution were exported and subsequently analyzed in Python for comparison with the CPMsi-derived clusters (Figure S4).

Clusters from the two methods were mapped by investing pixels overlaps (Table S2). Jaccard index was calculated (Figure S4 C) as the ratio of the number of shared pixels (intersection) to the total number of pixels assigned to either cluster (union). Values close to 1 indicate highly reproducible spatial partitions, whereas values close to 0 indicate little overlap. Most clusters were uniquely matched, indicating that each cluster identifies roughly the same region. To further investigate discrepancies between the two approaches, we examined cases in which a single *SCiLS Lab* cluster was divided into multiple CPMsi clusters. For example, SCiLS cluster 16 partitioned into four CPMsi clusters (Figure S4 C-D). Spectral similarity analysis of the corresponding consensus spectra revealed slightly lower similarity among the CPMsi-derived subclusters, indicating the presence of molecular heterogeneity that was not captured by the *SCiLS Lab* segmentation.

To identify the *m/z* features contributing to the reduced cosine similarity between CPMsi clusters 17 and 19, the presence and mean intensity of each feature detected in cluster 17 were evaluated within cluster 19 (Table S3). Table S2 ranks features according to their percentage presence in cluster 19 and highlights ions with lower occurrence frequencies. Several peaks prominent in the consensus spectrum of cluster 17 were present in fewer than 80% of spectra belonging to cluster 19, with m/z 872.6303 exhibiting the lowest prevalence (64%). Most remaining features were highly conserved (>80% presence), indicating that the two clusters share a common molecular profile but differ in the consistency of a limited number of discriminating ions. These findings support subtle molecular heterogeneity within the original *SCiLS Lab* cluster; however, the observed differences depend on the predefined consensus criteria, including the 90% peak presence threshold used to generate the consensus spectra (Figure S4 F).

**SI3. MS/MS Acquisition**

To demonstrate an alternative strategy for the annotation of CPMsi-derived molecular features when complementary LC-MS data are not available, we performed a dedicated MSI-MS/MS experiment. This experiment also serves to showcase the speed advantage of the CPMsi: molecular features can be detected and prioritized as soon as raw data are available from the instrument, enabling targeted MS/MS acquisition during the same measurement session. The resulting fragmentation spectra provide structural information that supports lipid annotation directly from the MSI experiment, facilitating biological interpretation without the need for additional analytical workflows.

We acquired MSI data from a brain tissue section in both positive- and negative-ion modes at a spatial resolution of 30 µm. The resulting datasets were processed independently using the CPMsi up to Module 3 (138,988 pixels processed in 10,23 min). Outputs were then integrated using the cluster-level integration framework described in Module 4, resulting in the identification of novel multimodal clusters (Figure S5). To characterize the molecular composition of the newly identified Cluster 26, *m/z* features were extracted according to their unique presence in the cluster. MS/MS measurements were performed on these peaks directly from the same tissue section. The acquired MS/MS spectra were searched against the *LIPID MAPS* glycerophospholipid database [4] using the experimentally determined precursor and product ion masses. To demonstration, we only report the MS/MS spectra of the precursor ion *m/z* 766.53 which was identified as PE(18:0_20:4) (Figure S6).

### Supplementary Figures

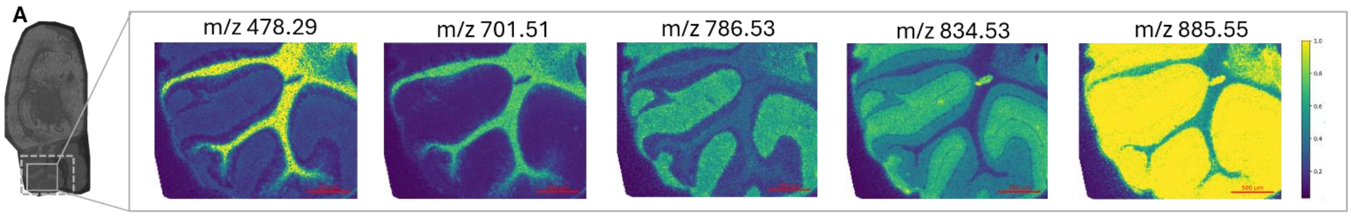


B

| Tolerance | Correlation $R^2$ | Run Time (s) | | CPMsi Speed Increase |
|---|---|---|---|---|
| | | CPMsi | MatchMS | |
| 0.05 | 0.901 | 622.40 | 10965.53 | 17.61x |
| 0.01 | 0.966 | 221.23 | 12077.39 | 54.6x |
| 0.02 | 0.972 | 11.37 | 13156.92 | 118x |

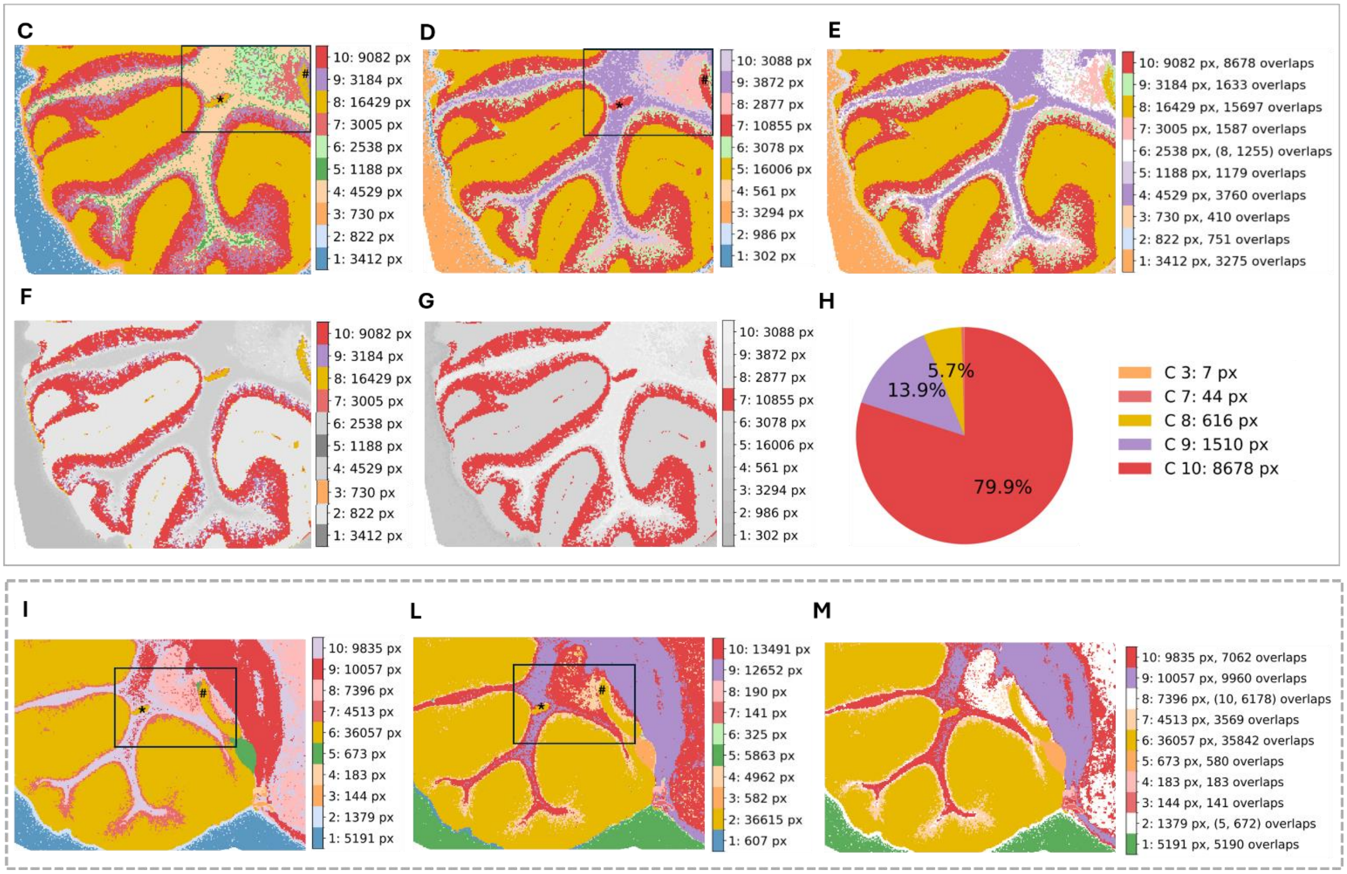


**Figure S1. Comparisons between CPMsi Pipeline and MatchMS Reference Pipeline.**

**(A)** Different ion spatial distributions within the 44,419 pixels sub-region (grey box). The dotted grey box represents an extended region of 75,428 pixels derived from enlarging the initial subregion **(B)** Run Time (s) and correlation scores between CPMsi and *matchms* with different tolerances. CPMsi is 118 times faster than *matchms* at the best tolerance 0.02. **(C)** Ten clusters obtained using the CPMsi. The black box highlights a detail of clustering disagreement between CPMsi Clusters and Reference Clusters **(D)** Ten clusters derived from hierarchical clustering of the reference cosine similarity matrix obtained from *matchms*. **(E)** Spatial overlap between the reference clusters **(D)** and CPMsi clusters **(C)**. The overlapping pixels from CPMsi clusters have been colored as pixels in reference clusters. **(F-G-H)** Example of overlaps: **(G)** cluster 7 from **(D)** partially overlaps with clusters 3-7-8-9-10 from **(C)** shown in **(F)**. The maximum pixels overlap (79.9%) is with cluster 10 **(H)**. **(I)** Ten clusters from CPMsi applied on the extended region. **(J)** Ten clusters from the *matchms* results applied on the extended region. **(K)** Spatial overlap between **(I)** and **(J)** with the **(I)** CPMsi clusters colored as **(J)** Reference clusters.

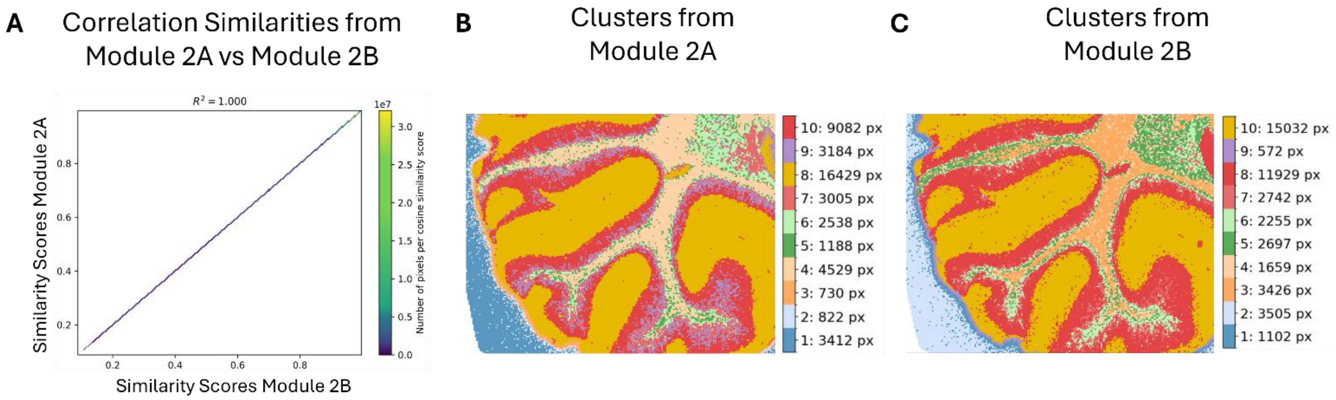


**Figure S2. Validation of Module 2B and 3B: comparison with Module 2A and 3A.**

**A)** Validation of Module 2B was conducted by performing Pearson Correlation between the cosine similarity matrices resulted from Module 2A and 2B; plot shows perfect correlation ($r^2$=1); **B)** Hierarchical Clustering from Module 3A; **C)** Hierarchical Clustering from Module 3B.

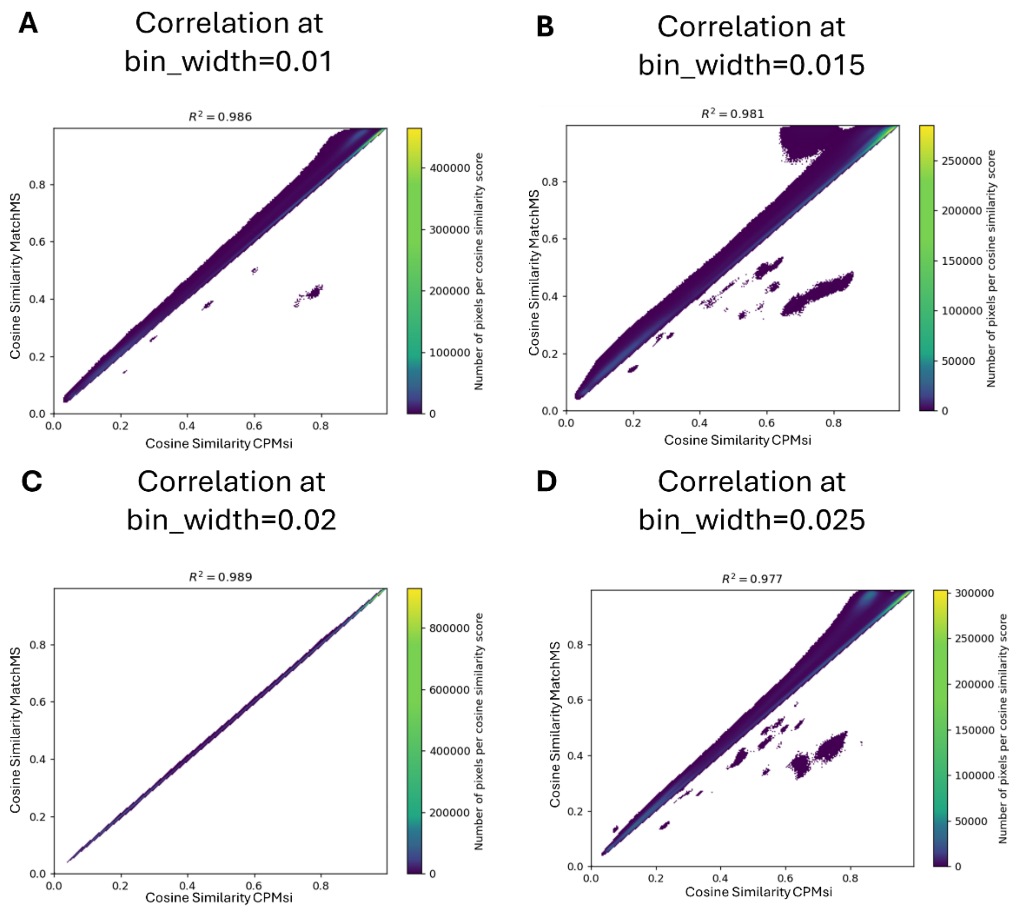


**E** Summary Data-driven Binning width

**Best: 0.02**

| Bin_width | $R^2$ | CPMsi Time (s) | MatchMS Time (s) |
|---|---|---|---|
| 0.01 | 0.985 | 41.99 | 131.27 |
| 0.015 | 0.980 | 25.56 | 138.80 |
| 0.02 | 0.989 | 19.07 | 145.35 |
| 0.025 | 0.977 | 18.62 | 151.90 |

**Figure S3. Density plot of cosine scores from CPMsi vs *matchms***

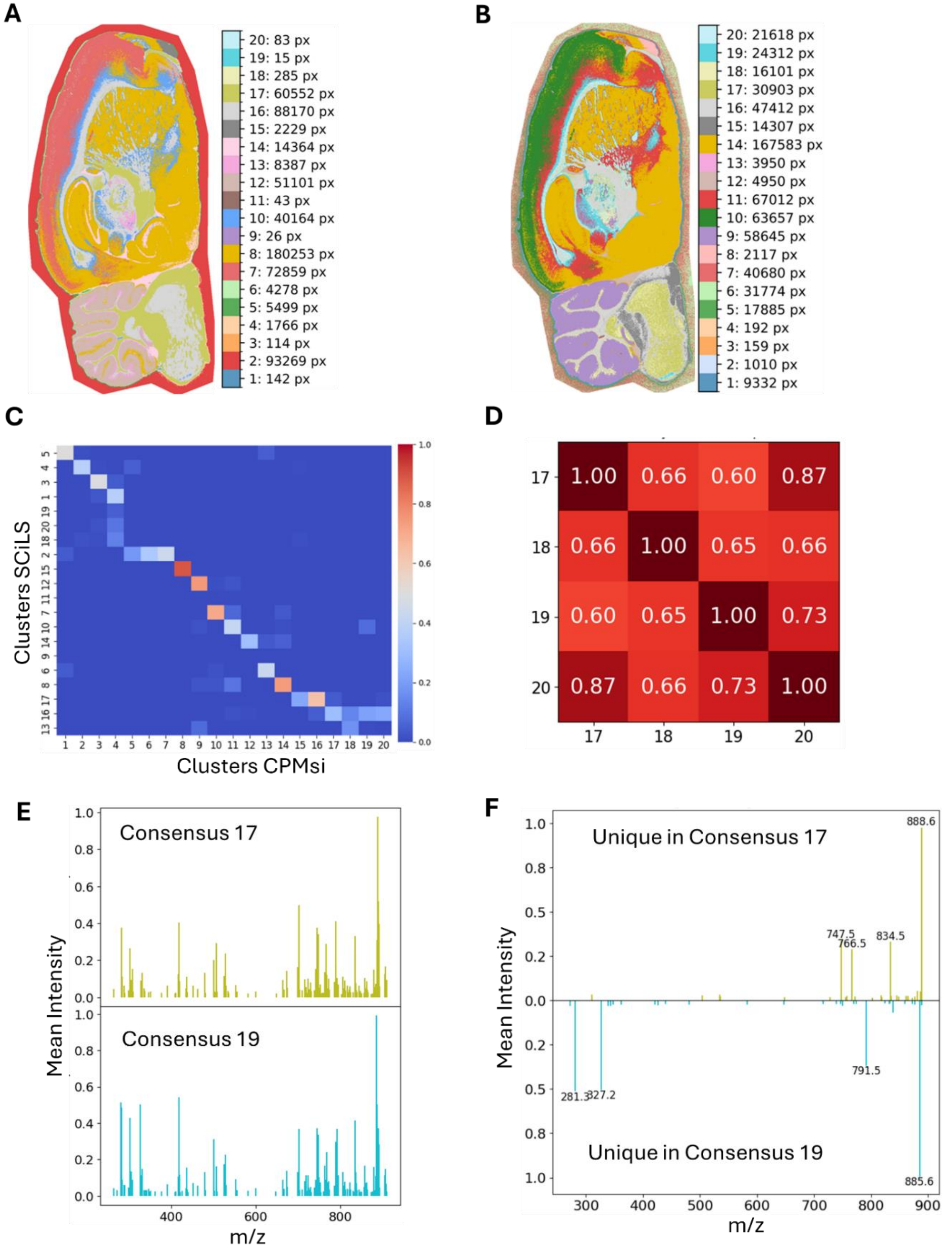


**Figure S4. Clusters mapping between *SCiLS Lab* segmentation map and CPMsi clusters.**

**A)** Hierarchical Clustering result from *SCiLS Lab*; **B)** Hierarchical Clustering result from CPMsi; **C)** Jaccard index between the two clustering; *SCiLS* Lab cluster 16 maps to CPMsi clusters 17-20; **D)** cosine similarity between CPMsi consensus spectra 17-20; **E)** consensus spectra generated by CPMsi from cluster 17 and 19 (cosine similarity= 0.60 as shown in D); **F)** *m/z* features that are uniquely present in each consensus (at the highest threshold of presence, i.e. peak present in more than 90% of pixels in the cluster).

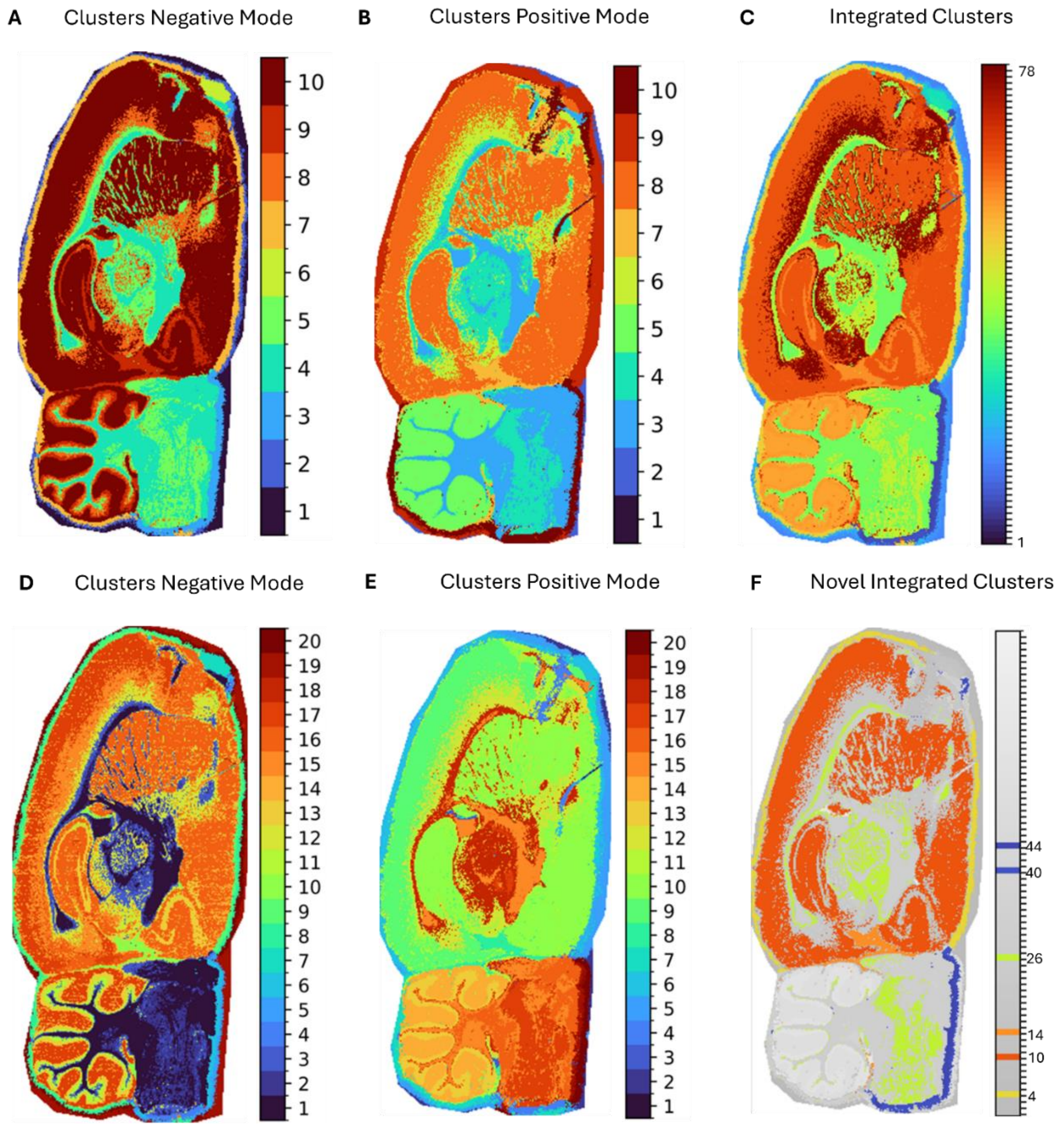

**Figure S5. Multimodal integration of positive- and negative-ion MSI datasets using CPMsi (A)** Spatial clustering of the negative-ion mode MSI dataset and **(B)** spatial clustering of the corresponding positive-ion mode MSI dataset; clusters are sorted and colored according to the similarity of their corresponding consensus spectra **(C)** Integrated clustering obtained by combining positive- and negative-ion mode with cluster-level integration strategy; **(D)** Higher-resolution clustering of the negative-ion mode dataset and **(E)** positive-ion mode dataset, revealing additional molecularly distinct tissue regions. **(F)** Novel clusters identified exclusively through multimodal integration, highlighting tissue regions that were not resolved in either ionization mode alone;

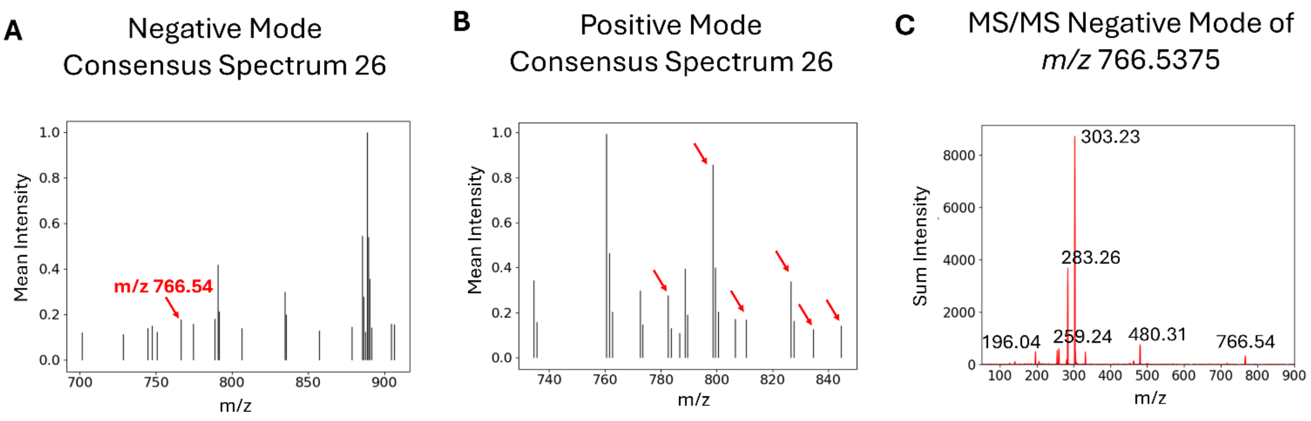


**D** LIPID MAPS Annotation

| Abbreviation | # of matches | m/z (delta) | Description |
|---|---|---|---|
| PE(18:0/20:4(5Z,8Z,11Z,14Z)) | 4 | | |
| | | 766.54 | Precursor ion [M-H]- |
| | | 500.28 | Loss of sn1 acyl chain as (RCH=C=O) from [M-H]- |
| | | 482.27 | Neutral loss of sn1 RCOOH group from [M-H]- |
| | | 480.31 (0.00) | Loss of sn2 acyl chain as (RCH=C=O) from [M-H]- |
| | | 462.30 | Neutral loss of sn2 RCOOH group from [M-H]- |
| | | 303.23 (0.00) | sn2 RCOO- ion |
| | | 283.26 (0.00) | sn1 RCOO- ion |
| | | 259.24 (0.00) | Loss of CO2 from sn2 RCOO- ion (PUFA) |
| | | 153.00 | Glycerol-3-phosphate ion with loss of H2O |
| | | 140.01 | Ethanolamine phosphate ion |
| | | 122.00 | Ethanolamine phosphate ion with loss of H2O |
| | | 96.97 | H2PO4- ion |
| | | 78.96 | PO3- ion |

**Figure S6. *m/z* features extraction followed by mid MS/MS acquisition run and *LIPID MAPS* Annotation. (A)** Negative-integrated consensus spectrum corresponding to integrated cluster 26; and **(B)** positive-integrated consensus spectrum corresponding to integrated cluster 26; red arrows denote uniquely present *m/z* features when compared to parent consensus **(C)** Negative-mode MS/MS spectrum of one the *m/z* features of integrated cluster 26 (precursor ion *m/z*

766.5375); **(D)** Spectral matching in *LIPID MAPS* identified the lipid as PE(18:0_20:4). Diagnostic fragment ions at *m/z* 303.23 and 283.26 correspond to arachidonate (20:4) and stearate (18:0) fatty acyl anions, respectively, while the ion at *m/z* 480.31 represents loss of the arachidonoyl chain. The observed fragmentation pattern is consistent with a phosphatidylethanolamine containing 18:0 and 20:4 acyl chains.

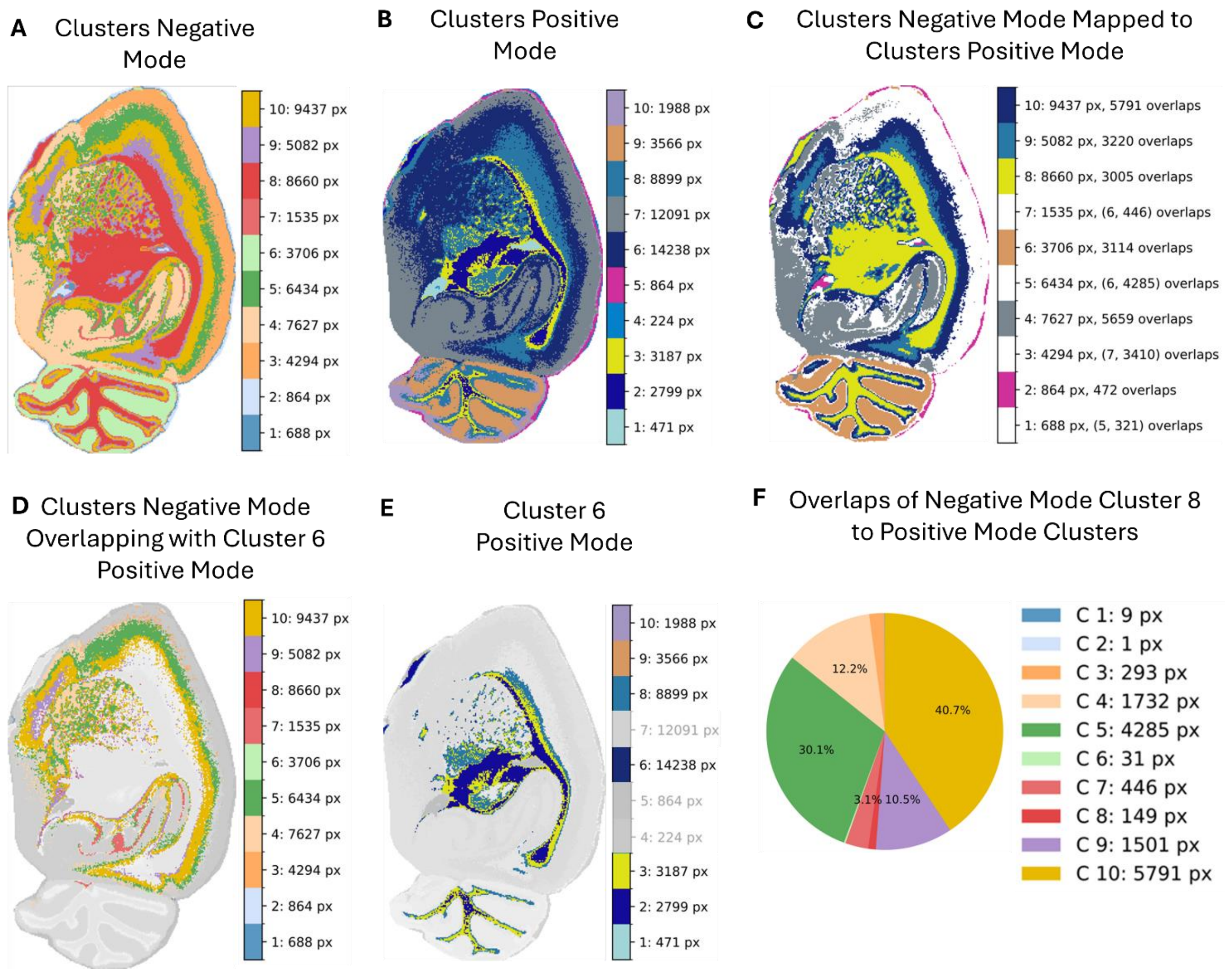


**Figure S7. Multi-omics application of the CPMsi Pipeline. (A)** Ten clusters of the Negative Mode dataset **(B)** Ten clusters of the Positive Mode dataset **(C)** Spatial overlap between (A) and (B) (Negative-ion mode clusters colored as Positive-ion mode clusters) **(D-E-F)** Example of overlaps:

**(D)** cluster 6 from (B) partially overlaps with clusters 1-2-3-6-8-9-10 from (A); **(E)** overlaps negative-ion mode cluster 8 with positive-ion mode clusters; **(F)** Percentage of overlaps of cluster 6 positive-ion mode with clusters of negative-ion mode.

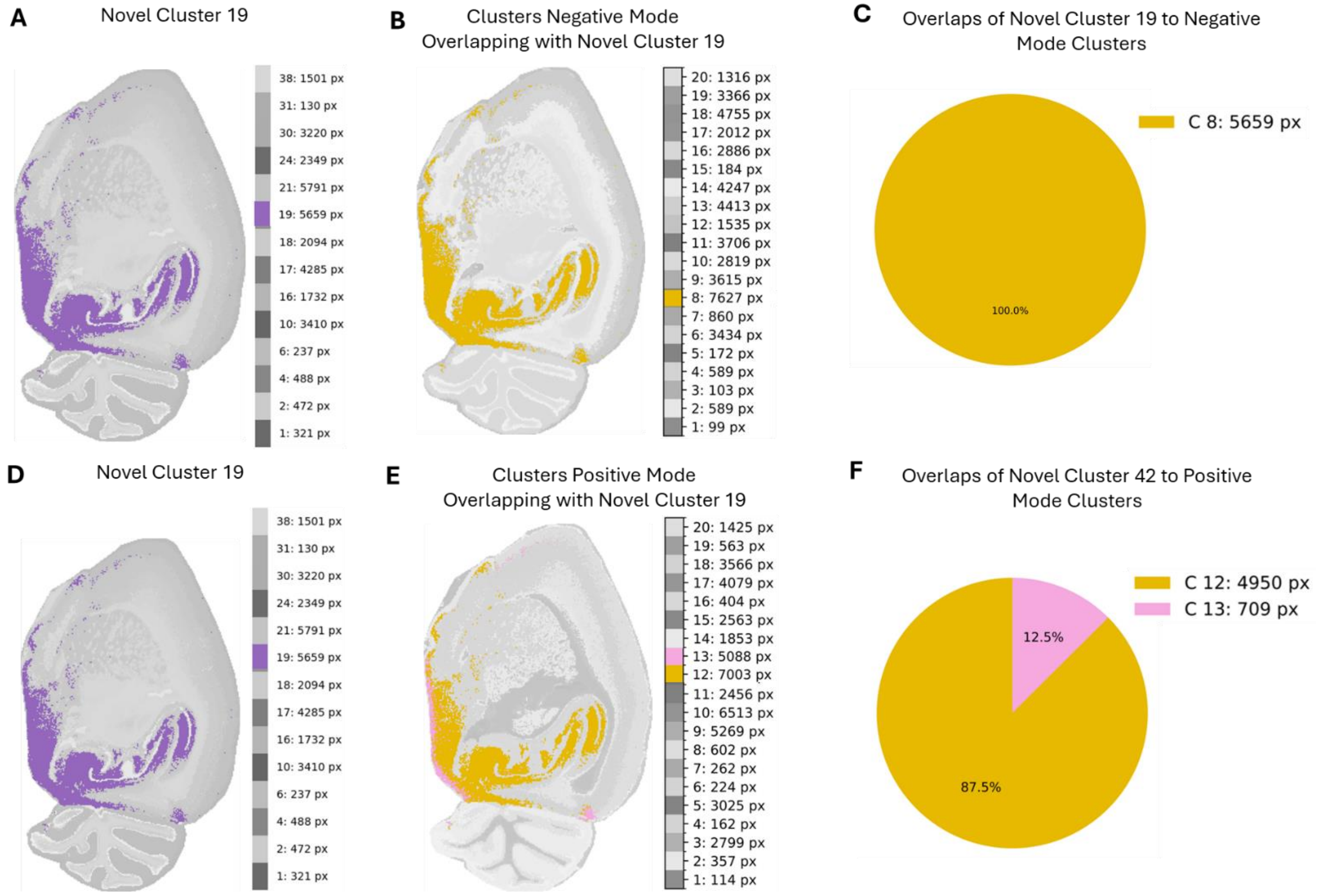


**Figure S8. Example of candidate novel cluster that is retrieved in the deeper-level clustering.** Candidate novel cluster 19 **(A)** is fully present in the 20-clustering negative mode **(B-C)** and positive mode **(E-F)**

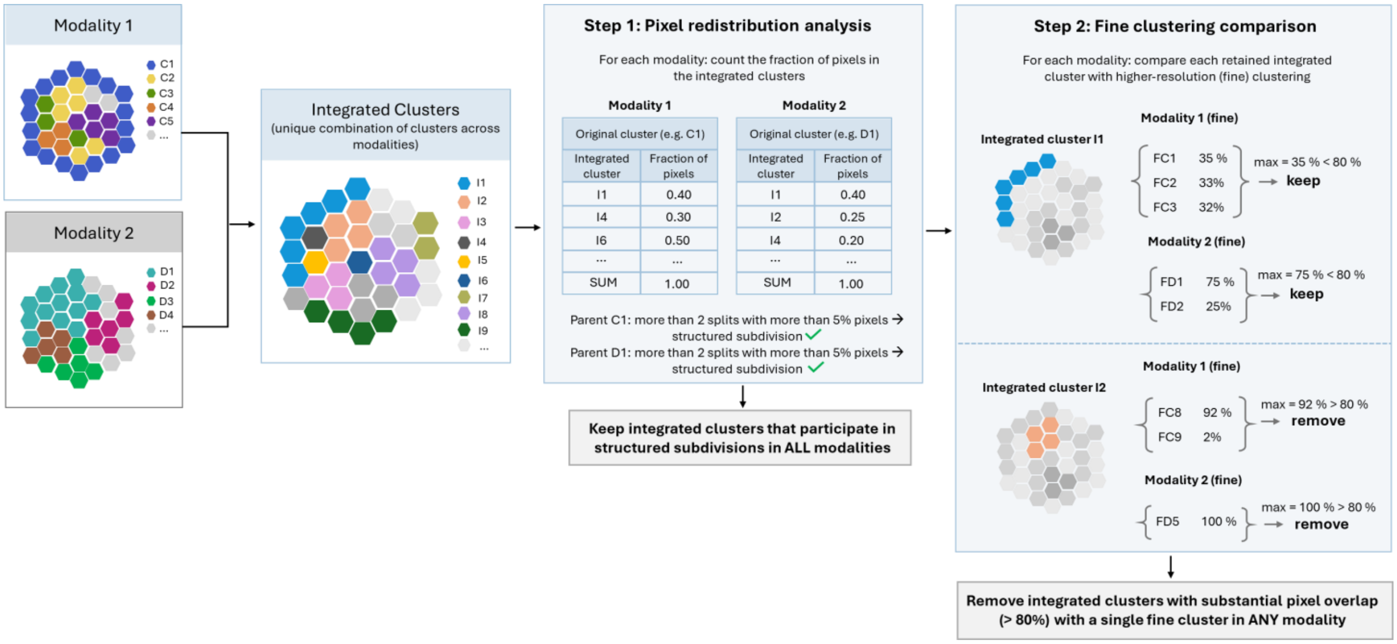


**Figure S9. Integration Strategy 1 for the identification of novel regions from multi-modal MSI data.** Independent clustering results from each modality are combined at matched spatial coordinates to generate integrated cluster labels. Candidate integrated clusters are first selected based on participation in structured subdivisions of parent clusters across all modalities. Retained clusters are subsequently compared with higher-resolution single-modality clustering results and discarded if they substantially overlap a single fine cluster. The remaining clusters are considered novel integrated regions emerging from multi-modal integration.

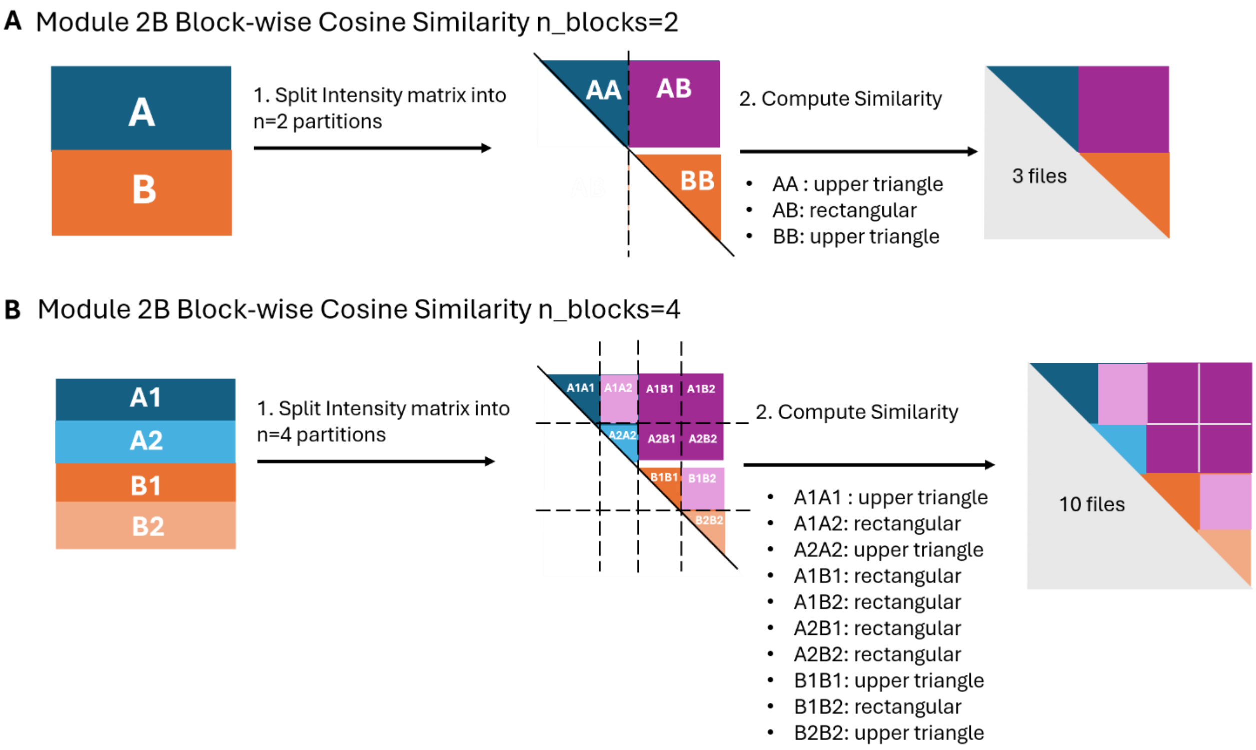


**Figure S10. Schematic illustration of the block-wise similarity calculation implemented in Module 2B.** Colored squares represent unique within-group and between-group pairwise comparisons, whereas the grey lower triangle indicates redundant symmetric comparisons that are not computed. **(A)** When the intensity matrix is divided into two partitions, the similarity calculation is performed in three blocks: two triangular blocks (AA and BB) and one rectangular block (AB). The results are stored in three separate files. When there are more partitions, this scheme is applied recursively to each partition. **(B)** For example, when the intensity matrix is divided into four partitions (A1, A2, B1, and B2), the upper-triangular block AA is further subdivided into two triangular blocks (A1A1 and A2A2) and one rectangular block (A1A2). Applying the same strategy across all partitions increases the total number of output files from 3 to 10.

| | 1 (n=1102) | 2 (n=3505) | 3 (n=3426) | 4 (n=1659) | 5 (n=2697) | 6 (n=2255) | 7 (n=2742) | 8 (n=11929) | 9 (n=572) | 10 (n=15032) |
|---|---|---|---|---|---|---|---|---|---|---|
| **1 (n=3412)** | 17 | 3395 | 0 | 0 | 0 | 0 | 0 | 0 | 0 | 0 |
| **2 (n=822)** | 712 | 110 | 0 | 0 | 0 | 0 | 0 | 0 | 0 | 0 |
| **3 (n=730)** | 373 | 0 | 0 | 0 | 0 | 0 | 0 | 1 | 356 | 0 |
| **4 (n=4529)** | 0 | 0 | 3400 | 412 | 716 | 1 | 0 | 0 | 0 | 0 |
| **5 (n=1188)** | 0 | 0 | 23 | 1018 | 144 | 3 | 0 | 0 | 0 | 0 |
| **6 (n=2538)** | 0 | 0 | 3 | 229 | 1691 | 615 | 0 | 0 | 0 | 0 |
| **7 (n=3005)** | 0 | 0 | 0 | 0 | 146 | 1599 | 1195 | 65 | 0 | 0 |
| **8 (n=16429)** | 0 | 0 | 0 | 0 | 0 | 1 | 10 | 1282 | 181 | 14955 |
| **9 (n=3184)** | 0 | 0 | 0 | 0 | 0 | 36 | 1456 | 1692 | 0 | 0 |
| **10 (n=9082)** | 0 | 0 | 0 | 0 | 0 | 0 | 81 | 8889 | 35 | 77 |

**Table S1. Table of overlapping pixels between Clusters Module 3A and Clusters Module 3B.** Module 3A clusters in the rows; Module 3B clusters in the columns.

| | **2 n=93269** | **5 n=5499** | **4 n=1766** | **1 n=142** | **3 n=114** | **20 n=83** | **19 n=15** | **18 n=285** | **14 n=14364** | **15 n=2229** | **12 n=51101** | **11 n=43** | **6 n=4278** | **9 n=26** | **7 n=72859** | **8 n=180253** | **17 n=60552** | **10 n=40164** | **13 n=8387** | **16 n=88170** |
|---|---|---|---|---|---|---|---|---|---|---|---|---|---|---|---|---|---|---|---|---|
| **1 n=51254** | 51245 | 3 | 0 | 5 | 0 | 0 | 0 | 1 | 0 | 0 | 0 | 0 | 0 | 0 | 0 | 0 | 0 | 0 | 0 | 0 |
| **2 n=35205** | 35013 | 16 | 157 | 6 | 10 | 2 | 1 | 0 | 0 | 0 | 0 | 0 | 0 | 0 | 0 | 0 | 0 | 0 | 0 | 0 |
| **3 n=6692** | 1245 | 4675 | 96 | 3 | 0 | 0 | 1 | 0 | 53 | 5 | 0 | 0 | 578 | 0 | 0 | 0 | 5 | 0 | 0 | 31 |
| **4 n=6179** | 5566 | 598 | 2 | 3 | 0 | 1 | 0 | 0 | 0 | 9 | 0 | 0 | 0 | 0 | 0 | 0 | 0 | 0 | 0 | 0 |
| **5 n=1675** | 179 | 3 | 1384 | 5 | 1 | 0 | 0 | 17 | 0 | 0 | 0 | 0 | 0 | 0 | 0 | 0 | 28 | 0 | 0 | 58 |
| **6 n=349** | 20 | 15 | 11 | 65 | 100 | 47 | 5 | 80 | 1 | 0 | 0 | 0 | 2 | 0 | 2 | 1 | 0 | 0 | 0 | 0 |
| **7 n=10** | 0 | 0 | 0 | 5 | 0 | 2 | 0 | 2 | 0 | 0 | 0 | 0 | 0 | 0 | 0 | 0 | 1 | 0 | 0 | 0 |
| **8 n=7264** | 0 | 4 | 0 | 11 | 0 | 0 | 1 | 2 | 6177 | 4 | 42 | 2 | 260 | 4 | 25 | 175 | 4 | 21 | 6 | 526 |
| **9 n=65279** | 0 | 0 | 0 | 1 | 0 | 0 | 0 | 0 | 7159 | 2 | 1 | 0 | 232 | 2 | 5 | 57690 | 0 | 185 | 0 | 2 |
| **10 n=2022** | 0 | 2 | 1 | 4 | 0 | 1 | 1 | 0 | 14 | 1971 | 0 | 0 | 12 | 0 | 0 | 11 | 0 | 0 | 0 | 5 |
| **11 n=55998** | 0 | 0 | 0 | 0 | 0 | 1 | 1 | 6 | 357 | 0 | 46796 | 15 | 24 | 0 | 1 | 4540 | 0 | 24 | 4186 | 47 |
| **12 n=3534** | 1 | 181 | 0 | 2 | 0 | 2 | 0 | 2 | 357 | 0 | 371 | 14 | 2439 | 3 | 108 | 7 | 0 | 5 | 5 | 37 |
| **13 n=53818** | 0 | 0 | 0 | 5 | 0 | 1 | 0 | 0 | 0 | 0 | 978 | 0 | 586 | 0 | 49737 | 2486 | 0 | 25 | 0 | 0 |
| **14 n=169723** | 0 | 2 | 0 | 1 | 0 | 0 | 1 | 0 | 227 | 223 | 2831 | 11 | 144 | 17 | 22978 | 115081 | 0 | 27267 | 65 | 875 |
| **15 n=20047** | 0 | 0 | 106 | 14 | 3 | 18 | 0 | 161 | 0 | 0 | 0 | 0 | 0 | 0 | 0 | 0 | 19741 | 0 | 0 | 4 |
| **16 n=41923** | 0 | 0 | 9 | 9 | 0 | 3 | 1 | 9 | 0 | 0 | 0 | 0 | 0 | 0 | 0 | 0 | 37106 | 0 | 0 | 4786 |
| **17 n=27619** | 0 | 0 | 0 | 0 | 0 | 0 | 0 | 0 | 16 | 9 | 3 | 1 | 0 | 0 | 2 | 258 | 0 | 11496 | 57 | 15777 |
| **18 n=15790** | 0 | 0 | 0 | 2 | 0 | 2 | 3 | 0 | 3 | 0 | 78 | 0 | 1 | 0 | 1 | 4 | 0 | 1134 | 3989 | 10573 |
| **19 n=24011** | 0 | 0 | 0 | 1 | 0 | 2 | 0 | 5 | 0 | 6 | 0 | 0 | 0 | 0 | 0 | 0 | 248 | 7 | 4 | 23738 |
| **20 n=35207** | 0 | 0 | 0 | 0 | 0 | 1 | 0 | 0 | 0 | 0 | 1 | 0 | 0 | 0 | 0 | 0 | 3419 | 0 | 75 | 31711 |

**Table S2. Table of mapped clusters between CPMsi and SCiLS.** CPMsi clusters in the rows; SCiLS clusters in the columns.

**Table S3. Presence of m/z features from consensus 17 in all pixels of cluster 19**

| mz | presence_pct | mean_intensity |
|---|---|---|
| 872.6303 | 63.9972 | 0.009843 |
| 755.5603 | 72.46627 | 0.01245 |
| 848.6403 | 77.32807 | 0.013448 |
| 771.5703 | 80.37595 | 0.013983 |
| 728.5303 | 81.48651 | 0.014484 |
| 757.5703 | 81.79911 | 0.015644 |
| 886.6003 | 82.05413 | 0.032818 |
| 534.3603 | 82.12817 | 0.016762 |
| 648.3803 | 83.76111 | 0.014973 |
| 876.6203 | 84.18476 | 0.016435 |
| 817.5803 | 84.29993 | 0.016219 |
| 835.6203 | 84.32873 | 0.016278 |
| 818.5703 | 84.7894 | 0.014917 |
| 504.3103 | 84.80174 | 0.017627 |
| 845.6503 | 84.87167 | 0.016558 |
| 864.6203 | 84.91691 | 0.016566 |
| 536.2503 | 85.1308 | 0.016316 |
| 864.5703 | 86.11385 | 0.0168 |
| 310.2803 | 87.92366 | 0.018395 |
| 802.5103 | 88.25683 | 0.015814 |
| 534.3203 | 88.28973 | 0.019376 |
| 864.6503 | 88.36788 | 0.018464 |
| 881.5203 | 88.91083 | 0.019328 |
| 861.6403 | 89.13294 | 0.018116 |
| 861.5903 | 89.42086 | 0.017617 |

**Algorithm 1. Block-wise GPU cosine similarity calculation.**

```
Input:
    X Intensity matrix (N × D)
    Rt Row tile size
    Ct Column tile size
Output:
    S Similarity block files for each partition

Start
    1. Normalize each row of X to unit L2 norm.
    2. Determine p=number of partitions according to memory availability
    3. Split X into partitions, for example when p=2:
        A = first half of rows
        B = second half of rows
    For each partition p:
        4. Compute S11 = A × A^T
            For each row tile Rt in X1
                For each column tile Ct in X1
                starting from the current row tile
                    G ← Rt × Ct^T
                    Store G in S11
                    If A ≠ B
                        Store G^T in the mirrored position
                    End If
                End For
            End For
            Save S11
        5. Compute S12 = A × B^T
            For each row tile Rt in A
                For each column tile Ct in B
                    G ← Rt × Ct^T
                    Store G in S12
                End For
            End For
            Save S12
        6. Compute S22 = B × B^T
            For each row tile Rt in X1
                For each column tile Ct in X1
                starting from the current row tile
                    G ← Rt × Ct^T
                    Store G in S11
                    If A ≠ B
                        Store G^T in the mirrored position
                    End If
                End For
            End For
            Save S22
    End for
```

### Supplementary Information References


1. Huber, F., Verhoeven, S., Meijer, C. & al., e., matchms - processing and similarity evaluation of mass spectrometry data. *J Open Source Softw* **5** (52) (2020).

2. Onoprishvili, T., Yuan, J., Petrov, K. & al., e., SimMS: a GPU-accelerated cosine similarity implementation for tandem mass spectrometry. *Bioinformatics* **41** (3) (2025).

3. Strohalm, M., Kavan, D., NováK, P., Volný, M. & Havlíček, H., mMass 3: A Cross-Platform Software Environment for Precise Analysis of Mass Spectrometric Data. *Anal Chem* **82** (11), 4648-4651 (2010).

4. Fahy, E. *et al.*, Update of the LIPID MAPS comprehensive classification system for lipids. *J Lipid Res* **50**, S9-14 (2009 ).
